# AI identifies potent inducers of breast cancer stem cell differentiation based on adversarial learning from gene expression data

**DOI:** 10.1101/2023.08.21.554075

**Authors:** Zhongxiao Li, Antonella Napolitano, Monica Fedele, Xin Gao, Francesco Napolitano

## Abstract

Cancer stem cells (CSCs) are a subpopulation of cancer cells within tumors that exhibit stem-like properties, and represent a potentially effective therapeutic target towards long-term remission by means of differentiation induction. By leveraging an Artificial Intelligence (AI) approach solely based on transcriptomics data, this study scored a large library of small molecules based on their predicted ability to induce differentiation in stem-like cells. In particular, a deep neural network model was trained using publicly available single-cell RNA-Seq data obtained from untreated human induced pluripotent stem cells at various differentiation stages and subsequently utilized to screen drug-induced gene expression profiles from the LINCS database. The challenge of adapting such different data domains was tackled by devising an adversarial learning approach that was able to effectively identify and remove domain-specific bias during the training phase. Experimental validation in MDA-MB-231 and MCF7 cells demonstrated the efficacy of 5 out of 6 tested molecules among those scored highest by the model. In particular, the efficacy of triptolide, OTS-167, quinacrine, granisetron, and A-443654 offer a potential avenue for targeted therapies against breast CSCs.

## Introduction

Cancer stem cells (CSCs) are a subpopulation of cancer cells within tumors that exhibit stem-like properties, including the ability to undergo self-renewal and asymmetric division giving rise to copies of themselves and the mature progeny of non-stem cells through differentiation. CSCs may mediate tumor metastasis and relapse, thus representing a potentially effective therapeutic target towards long-term remission by means of differentiation induction^1^. It has been noted that even a partial success of differentiation therapy could improve the prognosis of most patients by decades^2^. Differentiation therapy represents a paradigm case in Acute Myeloid Leukemia (AML), where terminal differentiation of CSCs has been shown to produce significant clinical benefits^3^. Although it has been proposed that such benefits in AML are not exclusively due to differentiation of CSCs, differentiation therapy still holds tremendous therapeutic hope, also for solid tumors^4–7^. In fact, CSCs have been identified in a broad spectrum of solid tumors^8^, including breast cancer^9^. It has also been demonstrated that, despite the fact that prolonged in vitro culturing is thought to result in loss of crucial stemness properties, established breast cancer cell lines possess a small fraction of self-renewing tumorigenic cells with the capacity to differentiate into phenotypically diverse progeny. Breast cancer stem cell (BCSC) content varies greatly among breast cancer cell lines and breast carcinomas^10,11^. Triple-negative breast cancers (TNBCs) contain large numbers of BCSCs while luminal breast tumors have lower stem cell contents^12,13^. Consistently, MCF7 luminal breast cancer cell line has a low percentage (0.7-1.4%) of BCSCs, while MDA-MB-231 TNBC cell line exhibits low or null CD24 expression and high percentage (more than 90%) of CD44+ cells^14^. BCSCs are able to undergo self-renewal, give rise to phenotypically diverse progeny and survive chemotherapy, thereby constituting an excellent model for CSCs^14^. Moreover, supporting evidence for a hierarchical CSC-based model of metastasis initiation has been provided through single-cell analysis of human metastatic breast cancer cells^15^. Stemness properties were also identified by analyzing transcriptomic data of breast cancer cells from patients^16^.

Given the potential of differentiation therapy and the evidence of CSCs in a broad spectrum of tumors, searching for small molecules that are able to target CSCs is an active area of research. For example, histone deacetylase inhibitors have been investigated for differentiation therapy in AML on the basis of their epigenetic effects^17^. In general, multiple methodologies have been proposed which leverage small molecule treatments to augment cell conversion^18^. These encompass numerous applications for cell reprogramming or transdifferentiation, including but not limited to, neurons^19^, endothelial cells^20^, pancreatic-like cells^21^, cardiomyocytes^22^, hepatocytes^23^, and other types of cells^24–26^. However, relatively poor understanding of differentiation mechanisms^2^ has prevented a systematic rational approach to the discovery of novel effective molecules. It is therefore unsurprising that, in the context of CSCs targeting, one of the major studies involved a high-throughput screening (HCS) approach, through which the ability of salinomycin in selectively killing BCSCs was discovered^27^. All these investigations underscore the potential of drug-enhanced cell type conversion, although they often require extensive experimentation and/or prior understanding of biological targets, making the screening of large small molecule libraries a remarkably difficult task. In contrast, computational methods can provide practical shortcuts to identify small sets of promising candidates for subsequent validations. We have previously introduced a general method for prioritizing small molecules in diverse cell conversion scenarios solely based on drug-induced transcriptional data, termed “DECCODE”^28^. The method’s efficacy was validated in a cell reprogramming protocol, showing promising results as a tool for differentiation studies as well. In particular, it was used to screen the LINCS^29^ database to search for stemness signatures among ∼20,000 drug-induced gene expression profiles.

While the DECCODE approach is based on classical statistics to match a single target profile, a large number of samples representing the desired transcriptional profile would allow for the application of more advanced machine learning models, which is the main methodological motivation for the present study (overviewed in **Figure 1**). Exploiting publicly available single-cell RNA-Seq (scRNA-Seq) data from human induced pluripotent stem cells (hiPSCs) labeled according to four differentiation stages, we devised an Artificial Intelligence (AI) approach to learn the corresponding expression patterns and subsequently prioritize drugs based on their ability to induce similar features. Such strategy poses the significant challenge of training an Artificial Neural Network from an scRNA-Seq dataset and using it to evaluate drug-induced profiles from the LINCS L1000 based collection, i.e., two completely different platforms and cellular contexts. We tackled the problem by developing “DREDDA” (“Drug Repositioning through Expression Data Domain Adaptation”), a domain-adaptive adversarial architecture that was able to learn and remove most of the domain-specific information from the two datasets while simultaneously solving the main task of identifying differentiation patterns. In particular, the technique allowed the model to learn domain-specific features (adversarial task) during the training phase and simultaneously avoid their use during differentiation stage classification (main task). Domain adaptation was first widely explored in visual recognition tasks, where it aims to apply visual recognition models trained in one domain (e.g., photos) to another domain (e.g., paintings)^30–32^. Along the same principles, DREDDA was designed to learn cell differentiation patterns from the scRNA-Seq dataset and use such acquired knowledge to predict the differentiation-induction ability of each drug from the LINCS collection. Finally, six of the most interesting hits from the resulting drug prioritization were experimentally validated, demonstrating the efficacy of 5 of them in reducing CSCs in MCF7 and MDA-MB-231 cell lines.

**Fig. 1.**
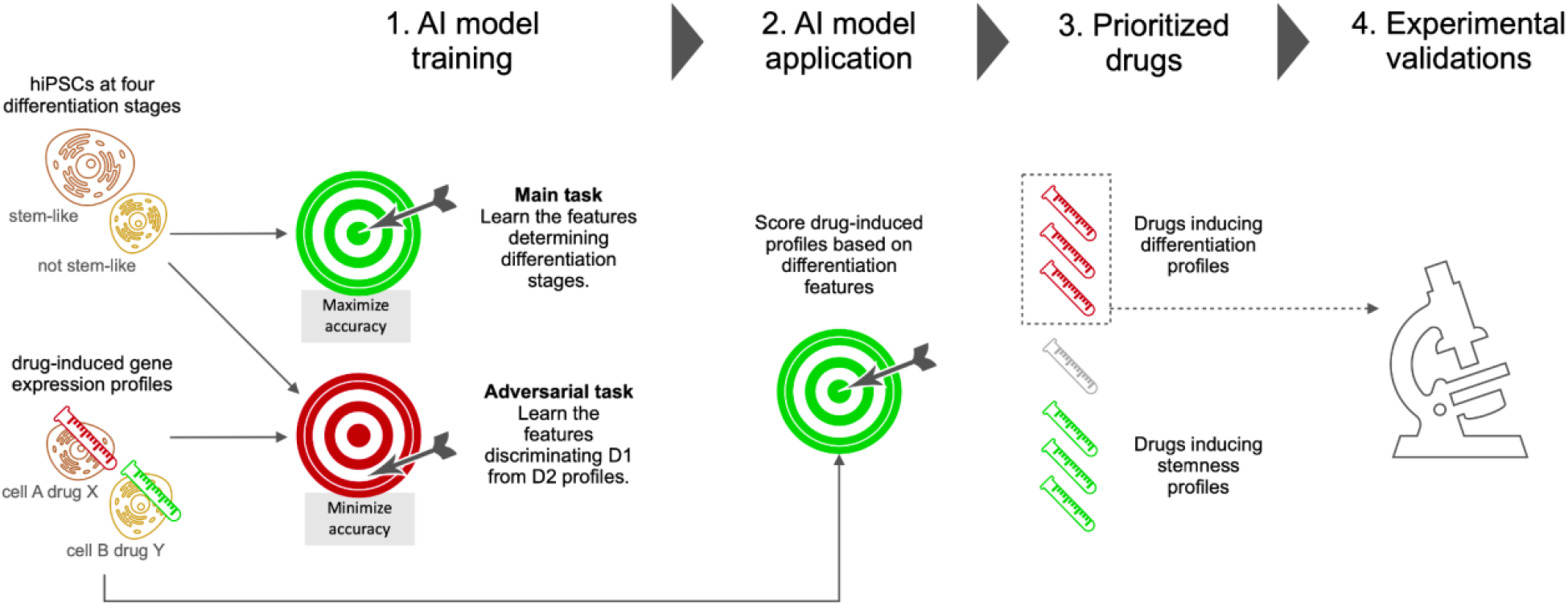
Overview of the study. Single-cell gene expression profiles of hiPSCs at various differentiation stages and drug-induced gene expression profiles were fed to an adversarial learning model, which simultaneously learned differentiation features to be used in subsequent predictions (main task) and dataset specific features to be avoided (adversarial task). The trained model was then used to score all the drug-induced profiles. A selection of 6 drugs among the top scoring ones was experimentally validated.

## Results

### Model development

With the aim of identifying BCSC differentiation inducing molecules, we designed an Artificial Intelligence (AI) approach completely based on transcriptional data (see Methods Section). The fundamental idea was to use a machine learning model in two steps: 1) learn transcriptional patterns that can discriminate stem cells from differentiated cells; 2) use the trained model to identify small molecules inducing similar patterns in treated cells. Towards this aim, we correspondingly exploited two different datasets: 1) a scRNA-seq dataset of human induced pluripotent stem cells (hiPSCs) including information about the differentiation stage of each cell; 2) a database of drug-induced transcriptional profiles obtained after the treatment of different cell lines. In particular, the scRNA-seq dataset we selected includes 18,787 hiPSCs obtained from WTC-CRISPRi^33^ cells. After sequencing, each cell was assigned one of four differentiation stages based on unsupervised clustering and biomarker analysis. As for the second dataset, we used drug-induced transcriptional profiles obtained from the LINCS database (GEO: GSE70138), including 107,404 differential gene expression profiles corresponding to the transcriptional responses of 41 cell lines to 1,768 different small molecules spanning different concentrations and time points^29^.

Since the model needs to be trained with the first dataset and provide predictions for the second one, the main challenge in its development was to effectively adapt the two domains, both of which are affected by biological and technical biases. The main source of biological bias came from the different cell types involved in both datasets. Although the cellular context represents an obviously relevant biological variable, it also acts as a severe limiting factor to the applicability of large drug-induced gene expression datasets. For this reason, methods treating cell type variability as biological bias have been proposed with the aim of maximizing drug prioritization performances from the available data^34^. Concerning technical biases, the two datasets were produced with remarkably different technologies, i.e. scRNA-Seq and L1000, the latter being specifically designed within the LINCS project. In order to reduce such sources of misleading signals, we devised an adversarial domain adaptation approach (see **Figure 2a** and Methods Section), in which a single deep learning model was trained to solve two competing tasks: 1) the main task, i.e. identifying the differentiation stage of each cell from the hiPSCs dataset; 2) the adversarial task, i.e. to discriminate between hiPSCs profiles and LINCS profiles (regardless of the treated cell line). In particular, the model was trained to maximize the performance of the main task and simultaneously minimize the performance of the second task. In this way, the extracted transcriptional features allowed the prediction of differentiation stages without relying on domain-specific information. During the training phase, the hiPSC dataset alone was used for the main task, while both datasets were used for the adversarial task. In particular, the training phase of DREDDA aimed for a steady increase of the main task classification performance and a steady decrease of the adversarial domain classification performance (**Figure 2b**). Indeed, the main task on the hiPSC dataset achieved 86.7% accuracy at the end of the training, significantly improving from the initial low performance. On the other hand, the adversarial task accuracy started at 100%, highlighting a severe dataset-dependent bias, but reached a ∼50% performance by the end of the training (**Figure 2b**), indicating near complete inability to distinguish between hiPSC and LINCS profiles. In other words, the information extracted by the model was sufficient to perform the main task, although largely irrelevant for the adversarial task. The internal representation of the data defined by the model after domain adaptation is visualized in **Figure 2c** together with a representation of the original data space. By comparing the two representations, it is evident how the clusters of cells belonging to each of the four differentiation stages appear significantly more separated after domain adaptation. On the other hand, the LINCS profiles, which mostly clustered together before adaptation, appear widely spread after adaptation, making them hardly separable from hiPSC profiles. We also quantified this effect by counting the percentage of hiPSC profiles falling in the 30 nearest neighbors of each LINCS profile before and after adaptation, showing a dramatic shift in the corresponding distributions (**Figure 2d**).

**Figure 2.**
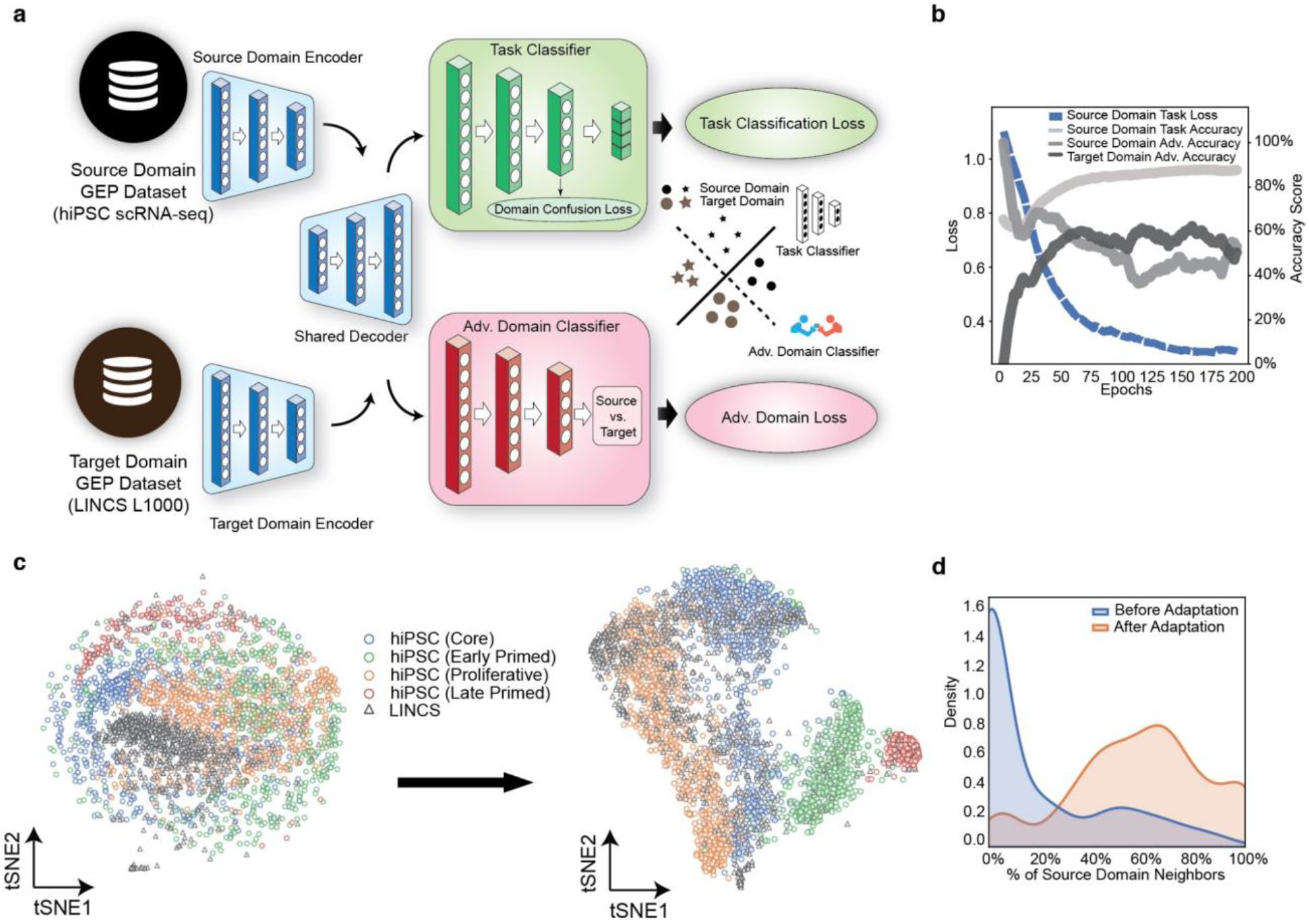
Model development. a) The DREDDA model architecture includes one encoder for each dataset and a shared decoder; the resulting profiles from the source domain are sent to the main task classifier (positively weighted in the overall loss function), while both source and target domain profiles are sent to the adversarial classifier (negatively weighted). b) During training, the main task accuracy increases, while the adversarial task accuracy decreases. c) Comparison between the embedding before (left) and after (right) domain adaptation shows that cells at the various differentiation stages tend to cluster together more, while LINCS drug-induced profiles tend to spread across the source domain. d) The neighborhood of untreated cell profiles tends to be more enriched for LINCS profiles after domain adaptation (orange) as compared to before (blue).

### Top hits validation and characterization

After training, the model was finally used to perform the main task on each of the LINCS profiles and thus predict the effectiveness of the corresponding treatment to induce the transcriptional features learned from the hiPSC dataset. In particular, we used the scores assigned by the model as a prioritization measure to rank all LINCS profiles. In order to validate the prioritization based on prior knowledge, we compared the DECCODE scores of the top 10 and bottom 10 drugs in the list. The DECCODE score is a gene expression data driven measure of stemness that we defined and validated in a previous study^28^. We observed low (high) DECCODE scores in the top- (bottom-) 30 drugs (**Figure 3a**), which appears coherent with differentiation (stemness) features. We also observed a general negative correlation between the DREDDA score and the DECCODE score (**Figure S1**).

**Figure 3.**
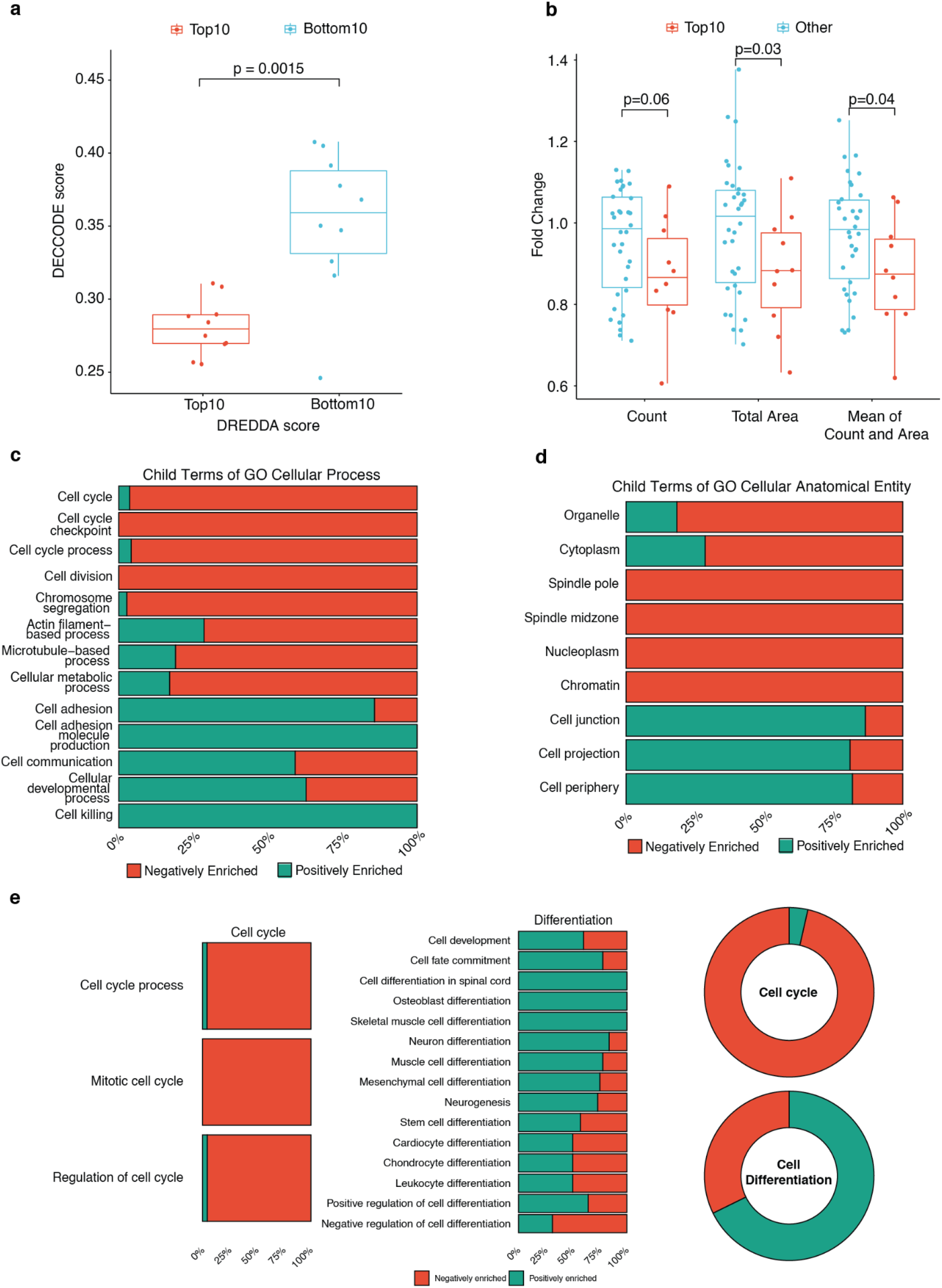
Validation and characterization of the top hits. a) Top (bottom) drugs as prioritized by DREDDA have low (high) DECCODE scores, which predict stemness features. b) Drugs previously tested for inducing stemness tend to be ranked lower by DREDDA based on experimental evidence including stem cells colony count and size. c-d) Summary of the positive and negative enrichments for pathways among the top levels of the “Biological Process” and “Cellular Component” Gene Ontology categories that are significantly dysregulated by the top 30 drugs. e) Same analysis as in c-d, but focused on the “Cell cycle” and “Differentiation” levels in the “Biological Process” category.

Apart from the validation of DREDDA’s prediction with the previous computational methods, we also specifically tested its consistency with previously published experimental results on a collection of 45 drugs, including 25 with high DECCODE scores and 20 with low DECCODE scores^28^. These drugs were tested for pluripotency induction in human inducible fibroblasts-like cells. After treatment, fold changes of cell colonies count and covered area were used to measure the efficacy of pluripotency induction. We sorted the list of 45 drugs according to their DREDDA scores and observed that the top-10 drugs generally induced lower colony count and colony-covered area in the DECCODE related experiments, thus experimentally confirming a general negative correlation between the two scores (**Figure 3b**).

Among the top-30 drugs prioritized by DREDDA (see **Table S1**). Many molecules in the set belong to chemotherapeutic agents in the class of kinase inhibitors, including CDK inhibitors, MELK inhibitors, JNK inhibitors. Other molecules specifically target DNA replication, including topoisomerase inhibitors and pyrimidine synthesis inhibitors. To investigate the common biological mechanisms affected by most drugs in this set, we performed a Drug Set Enrichment Analysis (DSEA)^35^ using the “Biological Process” and “Cellular component” categories of the Gene Ontology (GO) collection (**Tables S2** and **S3**). The most significant pathways enriched with a negative score included many that are associated with the cell cycle process (such as *cell cycle G2-M phase transition* - GO:0044839; *positive regulation of cyclin dependent protein kinase activity* - GO:1904031; *Telomerase RNA localization* - GO:0090672) and structures involved in it (including *nuclear envelope* - GO:0005635; *spindle pole* - GO:0000922; *centrosome* - GO:0090672). On the other hand, the most significant pathways with a positive enrichment score mostly concerned cell communication (e.g. *Regulation of calcium ion transmembrane transport* - GO:1903169; *Regulation of hormone levels* - GO:0010817; *Organic anion transport* - GO:0015711) or differentiation (*Pattern specification process* - GO:0007389; *Regionalization* - GO:0003002; *Photoreceptor cell differentiation* - GO:0046530). Next, in order to obtain a more high-level overview of the most recurrent cellular activities impacted by the drug set, we systematically investigated the up- and down-regulation of pathways falling within larger families of biological processes and cellular components. In particular, we quantified how many negatively and positively DSEA-enriched pathways fell below each one of the top terms in the GO hierarchy (**Figure 3c**). Notably, most pathways in the families of cell cycle (i.e. *cell cycle* - GO:0007049 itself; *cell cycle checkpoint* - GO:0000075; *cell cycle process* - GO:0022402) and cell division (i.e. *cell division* - GO:0051301 itself; *chromosome segregation* - GO:0007059; *actin filament-based process* - GO:0030029; *cellular metabolic process* - GO:0044237) were negatively enriched, suggesting a general inhibition of the cell cycle progression under the treatment of the top 30 drugs. In contrast, most pathways within families that are possibly related with cell differentiation (i.e. *cell adhesion* - GO:0007155; *cell communication* - GO:0007154; *cellular developmental process* - GO:0048869) were positively enriched. Consistently, the same analysis on top level pathways in the Cellular Component category showed that most cell cycle-related cellular structures were negatively enriched (e.g., *spindle pole* - GO:0000922; *nucleoplasm* - GO:0005654; and *chromatin* - GO:0000785), while those possibly related with differentiation through cell communication were positively enriched (*to cell junction* - GO:0030054; *cell projection* GO:0042995; cell periphery - GO:0071944) (**Figure 3d**). All such results were obtained by blindly investigating pathways and families of pathways without using any prior information. However, given the known desired effects that the drugs were prioritized for by DREDDA, we further investigated the enrichment of pathways below the *cell cycle* and *cell differentiation* levels in the GO hierarchy (**Figure 3e**). All of the three levels below *cell cycle* (i.e. *cell cycle process* - GO:0022402; *mitotic cell cycle* - GO:0000278; *regulation of cell cycle* - GO:0051726) were highly enriched by negatively regulated pathways. On the other hand, most levels below the *cell differentiation* term appeared positively enriched.

### Molecules inducing CSC differentiation

#### In vitro biological evaluation: Effects of the molecules on general breast cancer cell viability

Computational results were validated through in vitro experiments using the MCF7 (luminal triple-positive breast cancer) and MDA-MB-231 (mesenchymal-like triple-negative breast cancer) cell lines, chosen as models of breast cancer with low and high percentage of CSCs, respectively^36^. Six small molecules (**Table 1**), out of the top 50, were selected and tested for cell viability using increasing drug concentrations to establish the IC50 (**Figure S2**). According to the MTT findings, *triptolide* and *OTS-167* were highly cytotoxic in both cell lines with IC50 at nanomolar concentrations. *A-443654* showed similar IC50 as compared to *OTS-167* only on MCF7 cells, while it was less effective on the more staminal and therefore chemo-resistant MDA-MB-231 cell line. *Granisetron* and leflunomide were better tolerated by both cell lines, resulting in IC50 at micromolar concentrations. Finally, *quinacrine* showed an intermediate IC50 in the low micromolar for both cell lines. Based on these findings we chose the working concentrations to be used for each molecule in the following assays targeting CSCs. Two significantly different effective dosages were used for all the molecules, as detailed in **Table S4**.

**Table 1.**
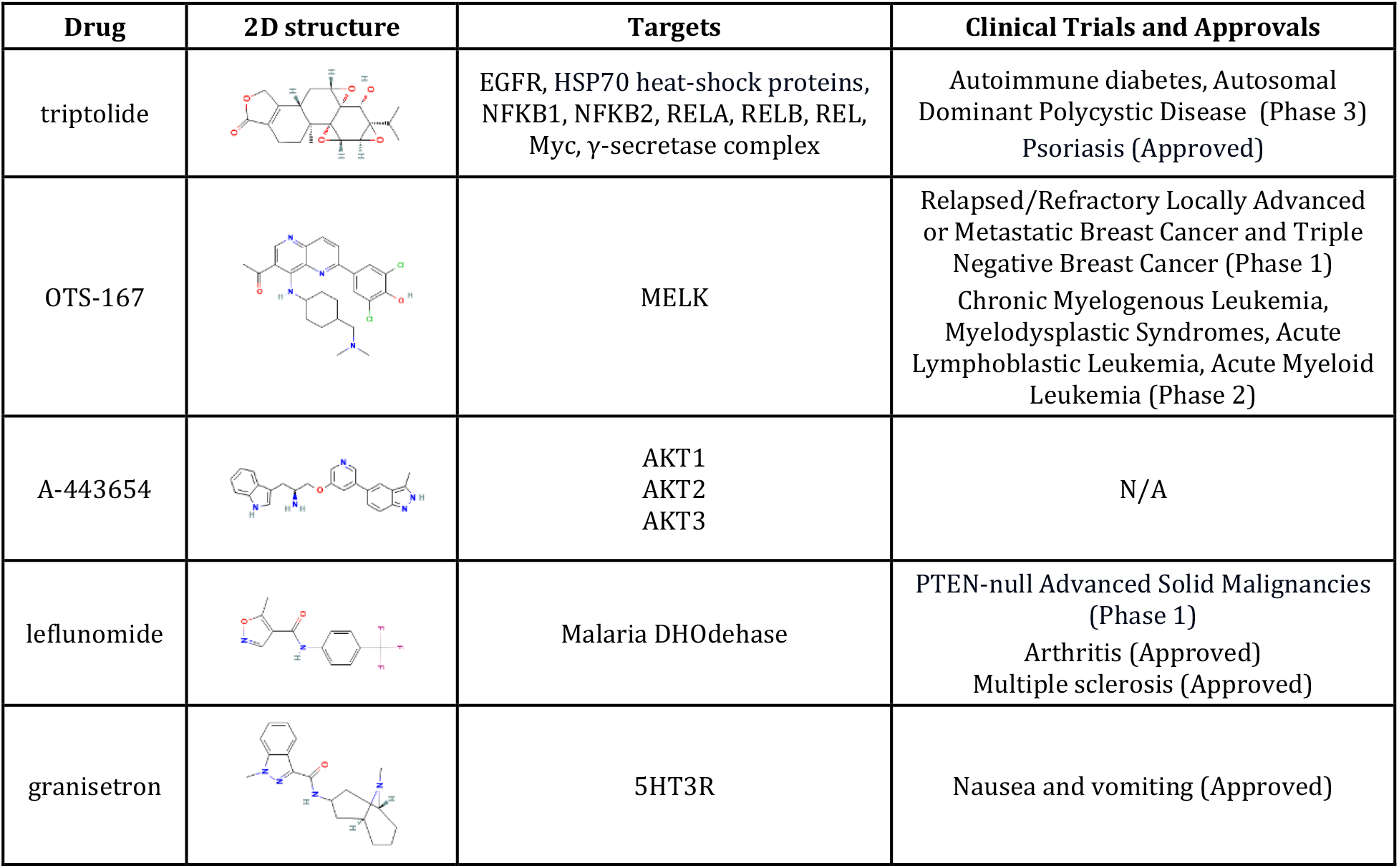
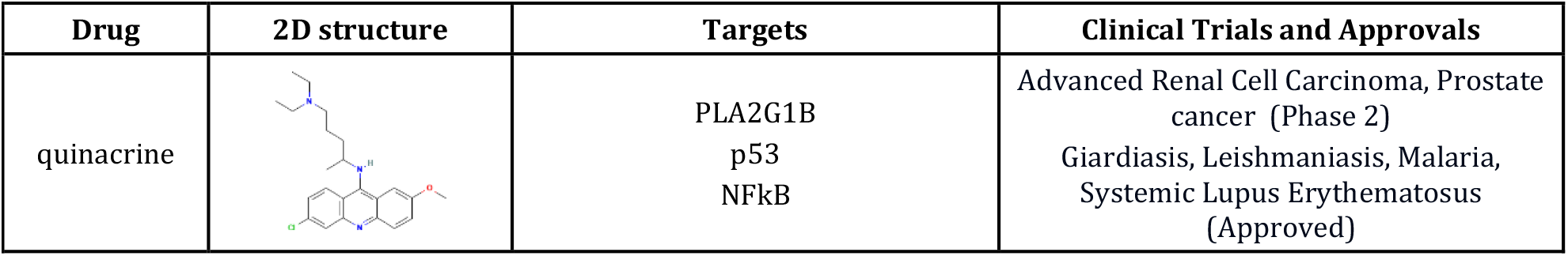
Small molecules selected for experimental validation from the top hits in the prioritization list.

#### Validation of molecule efficacy on breast cancer stem cells

To evaluate the effects of each drug on breast cancer stem cells (BCSCs), we first treated adherent cells for 24 hrs, then we washed out the drug and seeded the survived cells in stem cell medium on ultra-low attachment plates to let only BCSCs growing as mammospheres. The analysis of mammosphere forming efficiency (MFE), growth ability and self-renewal showed that three drugs, *Triptolide, OTS-167* and *Quinacrine*, effectively suppressed growth of BCSCs in both cell lines (**Figure 4a-b**). In more details, *Triptolide* decreased MFE of both MDA-MB-231 and MCF7 cells in a dose-dependent manner. It also reduced mammosphere growth ability and self-renewal for both cell lines by nearly 80% and 90% at the highest dose. *OTS-167* inhibited MFE and self-renewal activity of MDA-MB-231 cells in a dose-dependent manner, while their growth ability was significantly inhibited only at the highest dose. *OTS-167* also decreased MFE and growth of MCF7 cells in a dose-dependent manner, while self-renewal was highly reduced at both doses without significant differences between them. *Quinacrine* showed a significant effect on the reduction of MFE and self-renewal (dose-dependent only for MFE) of MCF7 cells, while its effect on MDA-MB-231 cells was only a significant reduction of mammosphere growing ability and a trend for a reduced MFE (**Figure 4b**). Other two drugs, *Granisetron* and *A-443654*, inhibited MFE, mammosphere growth and self-renewal of either MCF7 or MDA-MB-231, respectively (**Figure 4c-e**). For *Granisetron*, only the lower dose (300μM) showed a significant effect. Finally, *leflunomide* did not show any significant effect on BCSC availability and growth of both cell lines (**Figure 4e**).

**Figure 4.**
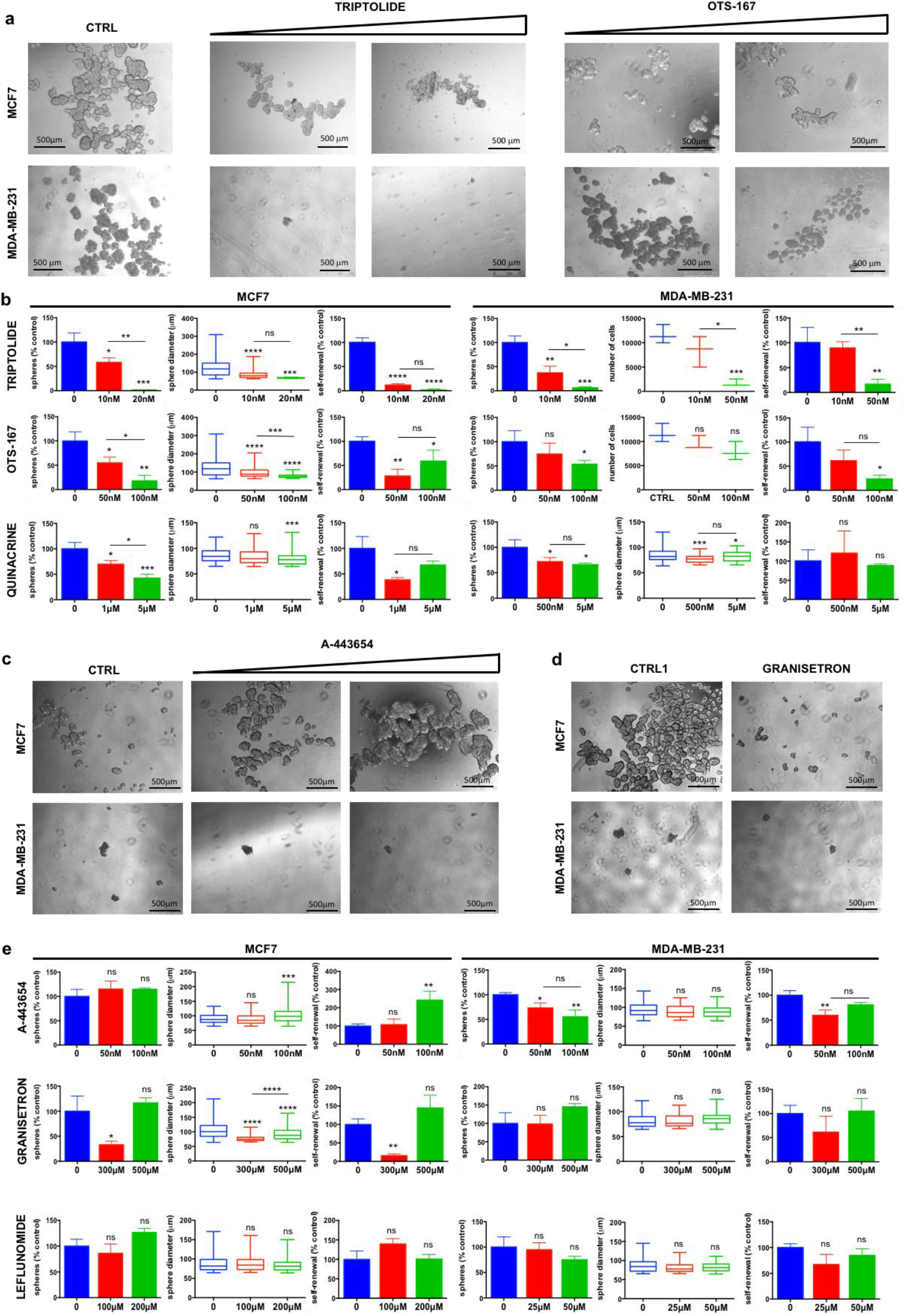
Mammosphere assay in drug-treated BC cells. (a) Representative images of MCF7 and MDA-MB-231 cells pre-treated for 24h with increasing doses of the indicated molecules and then cultured for 7 days as mammospheres in stem cell medium after the washing out of the drug. (b) Average number of mammospheres, their diameter and self-renewal capacity in three independent experiments. For MDA-MB-231 treated with triptolide and OTS-167, the number of single cells composing the mammospheres, as a measure of their growth, is reported instead of the mammosphere diameter. (c-d) Representative images of MCF7 and MDA-MB-231 cells pre-treated for 24h with increasing doses of each molecule and then cultured for 7 days as mammospheres in stem cell medium after the washing out of the drug. (d) For granisetron, only the lower dose (300μM) and its relative control (CTRL1) were shown. (e) Average number of mammospheres, their diameter and self-renewal capacity in three independent experiments. CTRL = DMSO 0.1%; CTRL1 = DMSO 0.6%; *, P < 0.05; **, P < 0.01; ***, P < 0.001; ****, P < 0.0001; ns, not significant.

#### Induction of BCSC differentiation

BCSCs are classically defined by CD44 (Cluster of Differentiation antigen-44) positive and low or absent levels of CD24 (Cluster of Differentiation antigen-24) expression (CD44^+^/CD24^−/low^) on their surface^37^ and recent clinical evidence has established that tumorigenic breast cancer cells with high expression of CD44 and low expression of CD24 are resistant to chemotherapy^38^. To evaluate whether the effect of the drugs on the availability and growth capacity of BCSCs was due to the induction of their differentiation, as predicted by the AI algorithm, we analyzed CD44 and CD24 expression by fluorescence-activating cell sorting (FACS), on adherent MDA-MB-231 and MCF7 cells treated for 24 hrs with each of the molecules that showed significant effects in the mammosphere assay. *Quinacrine* was found to be the most effective drug in inducing BCSC differentiation in both MCF7 and MDA-MB-231 cells, as assessed by the large dose-dependent decrease and simultaneous increase of the CD44^+^/CD24^-^ and CD44^-^/CD24^+^ subpopulations, respectively, in the MDA-MB-231 cells, and significant dose-dependent increase of the CD44-/CD24+ subpopulation in the MCF7 cells (**Figure 5a-b**). However, *Triptolide* and *OTS-167* also showed significant differentiating effects on both MCF7 and MDA-MB-231 cells (**Figure 5b**), while *Granisetron* and *A-443654* showed significant differentiating effects only on MCF7 or MDA-MB-231, respectively (**Figure 5c-d**), consistently with the mammosphere assay. These results confirm that the effects of the selected drugs on availability, growth capacity and self-renewal of BCSCs are due to the induction of their differentiation.

**Figure 5.**
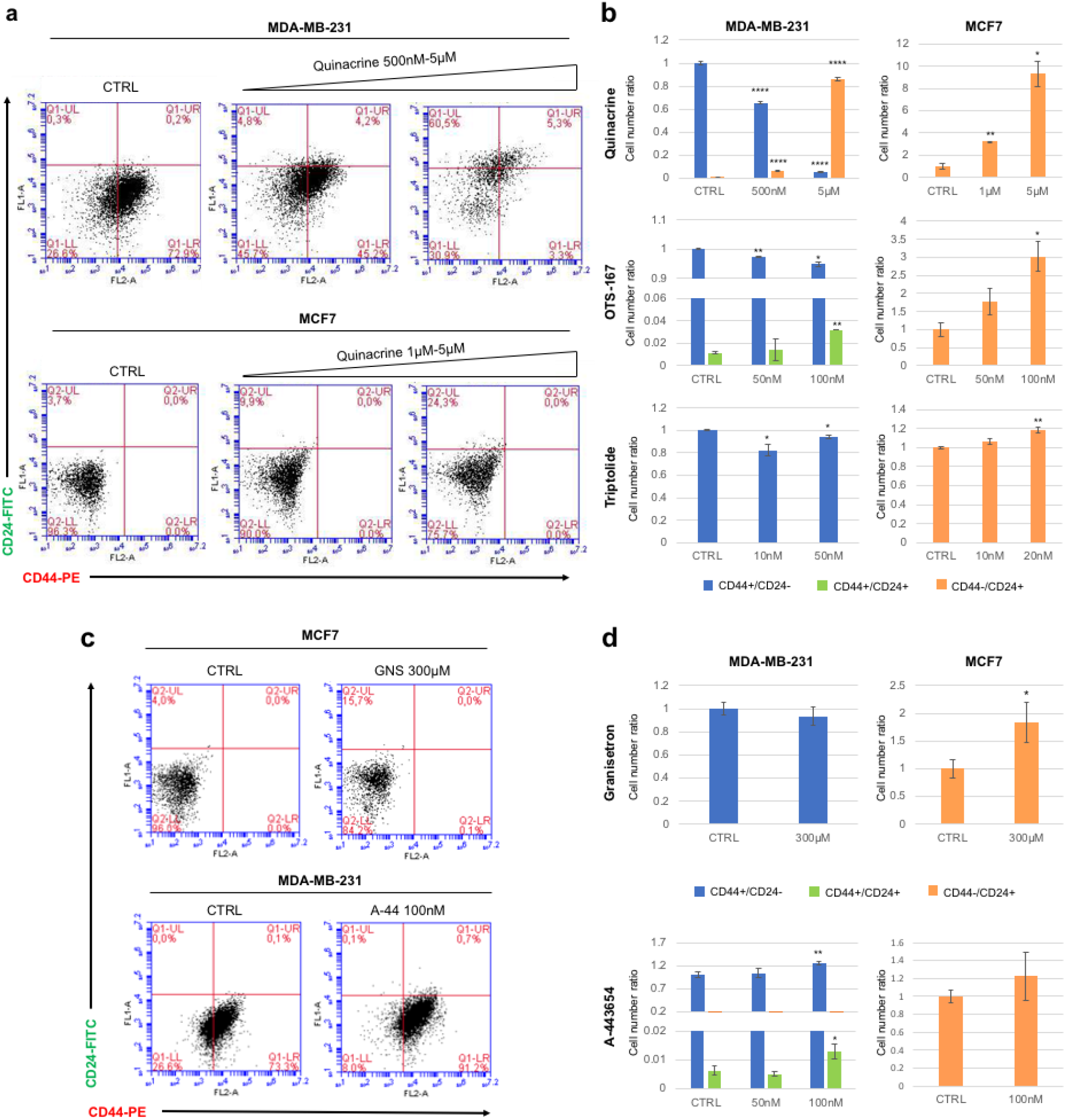
FACS profiling of CD44 and CD24 expression in MDA-MB-231 and MCF7 cells treated with quinacrine, triptolide, OTS-167, granisetron and A-443654. (a) Representative dot plots for quinacrine-treated cells. (b) The mean values +/- SE of the CD44+/CD24- (blue bars), CD44+/CD24+ (green bars) and CD44-/CD24+ (orange bars) subpopulations were reported as ratio relative to control (CTRL: DMSO 0.1%) for all treatments. (c) Representative dot plots of granisetron-treated MCF7 and A-443654-treated MDA-MB-231 cells. GNS, granisetron; A-44, A-443654. (d) The mean values +/- SE of of the CD44+/CD24- (blue bars), CD44+/CD24+ (green bars) and CD44-/CD24+ (orange bars) subpopulations were reported as ratio relative to control (CTRL: DMSO 0.1% for A-443654; DMSO 0.6% for granisetron) for all treatments. *, P < 0.05; **, P < 0.01; ****, P < 0.0001.

## Discussion

Breast cancer (BC) is a complex disease characterized by cellular heterogeneity among which the presence of cancer stem cells (CSCs) has been identified as a key factor contributing to tumor initiation, progression, and therapy resistance, thereby indicating an important therapeutic target. In this study, an artificial intelligence (AI) approach was employed to identify potential differentiating agents targeting breast cancer stem cells (BCSCs). The utilization of AI offered a powerful tool for screening a large library of compounds and identifying molecules with desired properties. Previous studies have demonstrated that drug-induced gene expression data, regardless of its known technical limitations and biological context dependency, can be effectively used to prioritize molecules facilitating cell type conversion based on a specific target expression profile. In this study, we showed how the same idea could be extended to an even more agnostic case, in which the target profile itself is not defined a priori. This was made possible by a machine learning approach that automatically extracted relevant transcriptional features from scRNA-Seq data. Indeed, from a computational perspective, single-cell transcriptomics proved to be an effective platform to obtain sizable datasets that are suitable for the training and testing of machine learning algorithms across diverse domains, despite the severe domain dataset biases involved. With the aim of ameliorating such bias, possibly relevant cell-specific features were likely removed during the training phase, which may represent the major drawback of this approach. Nonetheless, this strategy is necessary to deal with the limited availability of consistent drug-induced gene expression profile datasets, another significant challenge for this type of data driven discovery algorithms.

Following the AI-based screening, the study experimentally validated the efficacy of five out of six selected molecules, including triptolide, OTS-167, quinacrine, granisetron, and A-443654, in targeting BCSCs by inducing them to differentiate. Two commonly studied BC cell lines, MCF7 and MDA-MB-231, were used to assess the impact of these compounds on BCSCs, showing effective suppression of mammosphere forming efficiency, growth and self-renewal. The differentiation induction was confirmed by an altered protein expression associated with stemness and differentiation. Indeed, CD44+/CD24-subpopulation was reduced in MDA-MB-231, while CD24+ cells were increased in both MDA-MB-231 and MCF7 cells.

In several previous studies, *Triptolide*, a natural compound (diterpenoid tri-epoxide) derived from the Chinese herb *Tripterygium wilfordii*, has been found to exhibit potent anti-cancer properties, including anti-proliferative, anti-metastatic, and pro-apoptotic effects in various cancer types^39–42^. Some of these studies explored the potential of *triptolide* in targeting BCSCs, showing it inhibits multiple signaling pathways involved in self-renewal and maintenance of BCSCs, including c-Myc, Wnt/β-catenin and Notch pathways^43–45^. Consistent with our results, Li et al. demonstrated that *triptolide* inhibited self-renewal and induced a more differentiated phenotype in BCSCs, leading to reduced tumor growth and metastasis^46^.

Similarly, there have been studies investigating the role of *quinacrine*, a well-known antimalarial drug, in targeting CSCs^47^. Specifically, *quinacrine* treatment effectively inhibited cell proliferation, migration, invasion and representative metastasis markers of BCSCs^48,49^. However, no studies have so far shown a direct role of this agent on CSC properties in BC or other tumor models.

*OTS-167*, also known as *OTSSP167*, is an orally available MELK (Maternal embryonic leucine zipper kinase) inhibitor that is currently in phase I/II clinical trials for various tumors^50^. MELK induces carcinogenesis effects and is tightly associated with extended survival and accelerated proliferation of CSCs in various tumors, including glioblastoma and BC^51^. Consistently, MELK inhibition by *OTS-167* treatment significantly suppresses the proliferation and neurosphere formation in glioblastoma stem cells, in which MELK expression is enriched^52^. However, there is limited research specifically focused on the role of OTS-167 in BCSCs. Only Chung et al., in their pioneer study on the development of this compound, investigated its direct impact on BCSCs, demonstrating its efficacy in suppressing mammosphere formation and tumor growth in xenograft studies^53^. Here, we confirmed its efficacy in reducing BCSC availability, growth and self-renewal by mammosphere assays, also showing it induces their differentiation.

*Granisetron*, a selective serotonin receptor (5-HT3) antagonist, is primarily used as an antiemetic medication to prevent chemotherapy-induced nausea and vomiting^54^. While *granisetron* has been extensively studied in the context of managing chemotherapy-related symptoms, its specific role in directly targeting CSCs has not been investigated yet. Some studies have suggested that certain 5-HT3 receptor antagonists, including *granisetron*, may possess anti-CSC properties. These studies indicate that 5-HT3 receptor antagonists can modulate signaling pathways of CSCs^55,56^. Here for the first time we demonstrated that a specific dosage of granisetron (300μM) effectively inhibits BCSC properties in MCF7 cells and induce them to increase expression of the epithelial differentiation marker CD24, suggesting it acts as a differentiating agent in these cells.

*A-443654* is a small molecule inhibitor that primarily targets Akt kinases, a protein family involved in multiple cellular signaling pathways regulating cell survival, proliferation, and growth, the dysregulation of which has been implicated in various types of cancer^57^. Importantly, *A-443654* has been shown to inhibit glioblastoma stem-like cells with similar efficacy compared with traditionally cultured glioblastoma cell lines^58^, but there was still no research on its effects on BCSCs. In our study we showed it is effective in targeting MDA-MB-231-derived BCSCs with a weak differentiating effect.

Overall, the current study has important implications for the development of targeted therapies against BCSCs. The AI-driven identification of potential CSC differentiating agents expands the repertoire of molecules available for therapeutic interventions. Moreover, it highlights the power of AI in accelerating drug discovery and repurposing efforts, specifically in identifying molecules capable of targeting CSCs. By inducing CSC differentiation, these molecules hold the promise of reducing tumor heterogeneity, inhibiting self-renewal, and sensitizing CSCs to conventional therapies. Nonetheless, in order to understand the potential impact on clinical applications of our study, some important limitations must be taken into account: i) in vitro studies may not fully represent the complex and dynamic conditions of a living organism; ii) cell lines might not accurately recapitulate the complexity of the original tumor; iii) in vitro studies often have short experimental durations, which may not capture the long-term effects of the drug or the development of drug resistance in cancer stem cells over time.

Future perspectives of this work involve the translation of these findings into preclinical and clinical studies. In vivo models and patient-derived xenograft models should be employed to assess the therapeutic efficacy, safety, and pharmacokinetics of these compounds. Most of these molecules have already been tested on humans and considered safe, which constitutes an obvious advantage in terms of possible clinical translation. Additionally, further investigations are needed to elucidate the underlying molecular mechanisms by which these compounds induce CSC differentiation.

In conclusion, the integration of AI-driven screening and experimental validation provides a valuable approach to identify molecules capable of differentiating BCSCs. The findings of this study, including the efficacy of *triptolide, OTS-167, quinacrine, granisetron*, and *A-443654*, offer potential avenues for targeted therapies against BCSCs. This work lays the foundation for further research and development, bringing us closer to more effective and personalized treatments for BC patients.

## Methods

### Gene expression data

A single-cell RNA-seq dataset consisting of 18,787 WTC-CRISPRi^33^ human induced pluripotent stem cells (hiPSCs) was obtained from a previous study^33^, in which each cell was assigned one of four pluripotency stages (core pluripotent, proliferative, early primed for differentiation, late primed for differentiation). A zero-inflated negative binomial (ZINB) autoencoder model^59^ was used to normalize and denoise the dataset. Feature selection was performed by selecting the top 1000 genes showing highest mutual information (MI) between each feature (gene) and the differentiation stage labels. Concerning LINCS drug-induced profiles, the last release was obtained from GEO (ID:GSE70138). It includes 118,050 profiles obtained after treatment of 41 cell lines with 1,796 small-molecule compounds. In particular, population-control normalized differential profiles included in the level-5 distribution were used. Finally, only genes included both in the LINCS profiles and in the set selected from the hiPSC data were used to train the computational model. See “Supplementary methods” for additional details.

### Neural network model

The DREDDA architecture is a three-module composite deep neural network (DNN) consisting of (1) a domain-specific autoencoder (green part in **Figure 2a**); (2) the main task classifier (blue part in **Figure 2a**) (3); an adversarial domain classifier (red part in **Figure 2a**). The domain-specific autoencoder is an autoencoder with two independent encoders, one for each input dataset, and a shared decoder. The output of the decoder is sent to subsequent modules. The task classifier is a multi-layer perceptron providing classification probability for each of the four pluripotency stages. Its training is performed only on the source domain (hiPSC data) based on a cross-entropy loss function *L*_*cls*_. The adversarial domain classifier receives the same input as the main classifier, however it aims at providing a binary decision on whether such input comes from the source domain (hiPSCs) or the target domain (LINCS). Therefore, it is trained using both the source and target domain data with a binary cross-entropy loss function (*L*_*adv*_). Inspired by the Deep Domain Confusion (DDC) framework^32^, a third objective was introduced enforcing similarity of intermediate network values between source domain and target domain examples by minimizing a Maximum Mean Discrepancy (MMD)^60^ (*L*_*dc*_). The whole model was trained to simultaneously optimize the three mentioned functions according to the composite loss function: *L*_*cls*_ − *L*_*adv*_ + λ*L*_*dc*_. The details of the network architecture are listed in **Table S5**.

Model training was performed with a two-phases update per training step: phase 1 updates the minimization objective parameters (parameters of the source domain encoder, the target domain encoder, the shared decoder, and the task classifier), while the phase 2 updates the maximization objective parameters (the adversarial domain classifier). For each training step, an equal number of source domain and target domain examples were sampled. The model was implemented using PyTorch 1.8^61^ deep learning framework and requires an NVIDIA CUDA-capable GPU with ≥ 10 GB of memory. See “Supplementary methods” for additional details.

### Gene Set Enrichment Analysis of DREDDA-prioritized drugs

Drug-Set Enrichment Analysis (DSEA^35^), which is built on the classical Gene Set Enrichment Analysis (GSEA^62^), identifies pathways consistently dysregulated across a set of experimental conditions (such as drug treatments). DSEA was performed using the top-30 drugs prioritized by DREDDA as the foreground set and the bottom 30 as the background set on the LINCS level 5 dataset for the “C5 GO Biological Process” and “C5 GO Molecular Function” categories as obtained from the MsigDB v7.2^64^. The analysis was performed using the gep2pep R package^65^. Summaries for the positive and negative expression within families of pathways were obtained by counting the pathways with positive or negative enrichment scores that were found significant (p < 0.05) by the DSEA analysis and fell below a given level of the GO category. In this case, negative enrichments for GO terms starting with “Negative” (such as “Negative regulation of cell differentiation”) were counted as positive.

### Cell cultures and treatments

Human breast cancer MCF7 and MDA-MB-231 cell lines (American Type Culture Collection, ATCC, Manassas, VA) were grown in Dulbecco’s modified Eagle’s medium (DMEM, Sigma-Aldrich, Milan, Italy) and Roswell Park Memorial Institute-1640 medium (RPMI-1640, Sigma-Aldrich), respectively, supplemented with 10% heat-inactivated fetal bovine serum (FBS, Sigma-Aldrich) and 1% penicillin– streptomycin, in 95% air/5% CO2 atmosphere at 37 °C.

The drugs, IC50, and working concentrations for cell treatments are shown in Table S1. All drugs were dissolved in DMSO (stock solution) and further diluted in cell medium (working solution). Negative control was the solvent alone at its final concentration.

### Cell viability

Cell viability was assessed by CellTiter 96 AQueous One Solution Cell Proliferation Assay (Promega BioSciences Inc., San Luis Obispo, CA), according to the manufacturer’s instructions, 72 hours after cell treatment (5 × 10^3^ cells/well, 96-well plates). Each drug concentration was tested in at least three independent experiments for each molecule and each cell line.

### Mammosphere assay and CSC self-renewal

Cells were seeded at sub-confluence in 6-well plates. After 16 hrs they were treated with two different concentrations of each compound or DMSO (drug solvent) as negative control. At 24 hrs post-treatment, cells were washed out the serum and seeded (3000 cells/well) in 24-well low attachment plates with stem cell medium, consisting of 1% B27 (Invitrogen, Carlsbad, CA), 10ng/ml bFGF (Invitrogen), 20 ng/ml EGF (Sigma-Aldrich) in DMEM/F12 (SIGMA-Aldrich) supplemented with 1% penicillin/streptomycin) to form primary (P0) mammospheres. The number and diameter of P0 mammospheres were counted after a period of 7-14 days to estimate the number of treatment-resistant CSCs. Moreover, their diameter was measured to analyze their capacity to proliferate. Then, spheres were disaggregated, and single cells were replated (3000 cells/well) to form secondary (P1) mammospheres. P1 mammospheres were analyzed after further 7-14 days to calculate the CSC self-renewal (P1 mammospheres/P0 plated single cells x 100).

### FACS analysis

Adherent cells were treated for 24 hrs with two different concentrations of each compound or negative control (DMSO), then cells were washed with phosphate-buffered saline (PBS) and harvested with 0.05% trypsin/0.025% EDTA. Detached cells were washed and resuspended in PBS supplemented with 2% FBS, 0.2% sodium azide (Staining buffer). Combinations of fluorochrome-conjugated monoclonal antibodies against human CD44 (PE; Santa Cruz cat. # SC-18849-PE) and CD24 (FITC; Beckton Dickinson cat. # BD555427) were added to the cell suspension (1 x 10^6^ cells) and incubated at 4°C in the dark for 30 min. The labeled cells were washed in Staining buffer and then analyzed on a BD Accuri Flow Cytometer (BD).

### Statistics

Ordinary one-way analysis of variance (ANOVA) followed by Tukey’s multiple comparisons test, through GraphPad Prism 6 software, was applied for comparison of groups of experimental data. Values analyzed are the average +/- SD of at least three independent experiments.

## Data and code availability

Code for the DREDDA model and relevant datasets for reproduction of the results are available at https://github.com/lzx325/DREDDA.

## Bibliography

1. Wicha, M. S. Targeting self-renewal, an Achilles’ heel of cancer stem cells. Nat. Med. 20, 14–15 (2014).

2. Cruz, F. D. & Matushansky, I. Solid Tumor Differentiation Therapy – Is It Possible? Oncotarget 3, 559–567 (2012).

3. Sachs, L. The control of hematopoiesis and leukemia: from basic biology to the clinic. Proc. Natl. Acad. Sci. 93, 4742–4749 (1996).

4. de Thé, H. Differentiation therapy revisited. Nat. Rev. Cancer 18, 117–127 (2018).

5. Jiang, W., Peng, J., Zhang, Y., Cho, W. C. S. & Jin, K. The Implications of Cancer Stem Cells for Cancer Therapy. Int. J. Mol. Sci. 13, 16636–16657 (2012).

6. Li, Y., Atkinson, K. & Zhang, T. Combination of chemotherapy and cancer stem cell targeting agents: Preclinical and clinical studies. Cancer Lett. 396, 103–109 (2017).

7. Yang, C., Jin, K., Tong, Y. & Cho, W. C. Therapeutic potential of cancer stem cells. Med. Oncol. Northwood Lond. Engl. 32, 619 (2015).

8. Chen, K., Huang, Y. & Chen, J. Understanding and targeting cancer stem cells: therapeutic implications and challenges. Acta Pharmacol. Sin. 34, 732–740 (2013).

9. Al-Hajj, M., Wicha, M. S., Benito-Hernandez, A., Morrison, S. J. & Clarke, M. F. Prospective identification of tumorigenic breast cancer cells. Proc. Natl. Acad. Sci. 100, 3983–3988 (2003).

10. Pece, S. et al. Biological and molecular heterogeneity of breast cancers correlates with their cancer stem cell content. Cell 140, 62–73 (2010).

11. Charafe-Jauffret, E. et al. Breast cancer cell lines contain functional cancer stem cells with metastatic capacity and a distinct molecular signature. Cancer Res. 69, 1302–1313 (2009).

12. Honeth, G. et al. The CD44+/CD24-phenotype is enriched in basal-like breast tumors. Breast Cancer Res. BCR 10, R53 (2008).

13. Park, S. Y. et al. Heterogeneity for stem cell-related markers according to tumor subtype and histologic stage in breast cancer. Clin. Cancer Res. Off. J. Am. Assoc. Cancer Res. 16, 876–887 (2010).

14. Fillmore, C. M. & Kuperwasser, C. Human breast cancer cell lines contain stem-like cells that self-renew, give rise to phenotypically diverse progeny and survive chemotherapy. Breast Cancer Res. BCR 10, R25 (2008).

15. Lawson, D. A. et al. Single-cell analysis reveals a stem-cell program in human metastatic breast cancer cells. Nature 526, 131–135 (2015).

16. Chung, W. et al. Single-cell RNA-seq enables comprehensive tumour and immune cell profiling in primary breast cancer. Nat. Commun. 8, 15081 (2017).

17. Bots, M. et al. Differentiation therapy for the treatment of t(8;21) acute myeloid leukemia using histone deacetylase inhibitors. Blood 123, 1341–1352 (2014).

18. Federation, A. J., Bradner, J. E. & Meissner, A. The use of small molecules in somatic-cell reprogramming. Trends Cell Biol. 24, 179–187 (2014).

19. Ladewig, J. et al. Small molecules enable highly efficient neuronal conversion of human fibroblasts. Nat. Methods 9, 575–578 (2012).

20. Sayed, N. et al. Transdifferentiation of human fibroblasts to endothelial cells: role of innate immunity. Circulation 131, 300–309 (2015).

21. Zhu, S. et al. Reprogramming of human primary somatic cells by OCT4 and chemical compounds. Cell Stem Cell 7, 651–655 (2010).

22. Cao, N. et al. Conversion of human fibroblasts into functional cardiomyocytes by small molecules. Science 352, 1216–1220 (2016).

23. Lim, K. T. et al. Small Molecules Facilitate Single Factor-Mediated Hepatic Reprogramming. Cell Rep. 15, 814–829 (2016).

24. Cheng, L. et al. Direct conversion of astrocytes into neuronal cells by drug cocktail. Cell Res. 25, 1269–1272 (2015).

25. Li, J. et al. Artemisinins Target GABAA Receptor Signaling and Impair α Cell Identity. Cell 168, 86-100.e15 (2017).

26. Wang, Y. et al. Conversion of Human Gastric Epithelial Cells to Multipotent Endodermal Progenitors using Defined Small Molecules. Cell Stem Cell 19, 449–461 (2016).

27. Gupta, P. B. et al. Identification of selective inhibitors of cancer stem cells by high-throughput screening. Cell 138, 645–659 (2009).

28. Napolitano, F. et al. Automatic identification of small molecules that promote cell conversion and reprogramming. Stem Cell Rep. 16, 1381–1390 (2021).

29. Keenan, A. B. et al. The Library of Integrated Network-Based Cellular Signatures NIH Program: System-Level Cataloging of Human Cells Response to Perturbations. Cell Syst. 6, 13–24 (2018).

30. Csurka, G. A Comprehensive Survey on Domain Adaptation for Visual Applications. in Domain Adaptation in Computer Vision Applications (ed. Csurka, G.) 1–35 (Springer International Publishing, 2017). doi:10.1007/978-3-319-58347-1_1.

31. Ganin, Y. & Lempitsky, V. Unsupervised domain adaptation by backpropagation. in Proceedings of the 32nd International Conference on International Conference on Machine Learning -Volume 37 1180–1189 (JMLR.org, 2015).

32. Tzeng, E., Hoffman, J., Zhang, N., Saenko, K. & Darrell, T. Deep Domain Confusion: Maximizing for Domain Invariance. ArXiv (2014).

33. Nguyen, Q. H. et al. Single-cell RNA-seq of human induced pluripotent stem cells reveals cellular heterogeneity and cell state transitions between subpopulations. Genome Res. 28, 1053–1066 (2018).

34. Iorio, F. et al. Discovery of drug mode of action and drug repositioning from transcriptional responses. Proc. Natl. Acad. Sci. 107, 14621–14626 (2010).

35. Napolitano, F., Sirci, F., Carrella, D. & Di Bernardo, D. Drug-Set Enrichment Analysis: A Novel Tool to Investigate Drug Mode of Action. Bioinformatics 32, 235–241 (2016).

36. Vazquez-Santillan, K. et al. NF-kappa?-inducing kinase regulates stem cell phenotype in breast cancer. Sci. Rep. 6, 37340 (2016).

37. Ponti, D. et al. Isolation and in vitro propagation of tumorigenic breast cancer cells with stem/progenitor cell properties. Cancer Res. 65, 5506–5511 (2005).

38. Li, X. et al. Intrinsic resistance of tumorigenic breast cancer cells to chemotherapy. J. Natl. Cancer Inst. 100, 672–679 (2008).

39. Liang, X. et al. Inhibition of RNA polymerase III transcription by Triptolide attenuates colorectal tumorigenesis. J. Exp. Clin. Cancer Res. CR 38, 217 (2019).

40. Zhang, Y.-Q. et al. Galactosylated chitosan triptolide nanoparticles for overcoming hepatocellular carcinoma: Enhanced therapeutic efficacy, low toxicity, and validated network regulatory mechanisms. Nanomedicine Nanotechnol. Biol. Med. 15, 86–97 (2019).

41. McGinn, O. et al. Inhibition of hypoxic response decreases stemness and reduces tumorigenic signaling due to impaired assembly of HIF1 transcription complex in pancreatic cancer. Sci. Rep. 7, 7872 (2017).

42. Han, Y. et al. Triptolide Inhibits the AR Signaling Pathway to Suppress the Proliferation of Enzalutamide Resistant Prostate Cancer Cells. Theranostics 7, 1914–1927 (2017).

43. Sarkar, T. R. et al. GD3 synthase regulates epithelial–mesenchymal transition and metastasis in breast cancer. Oncogene 34, 2958–2967 (2015).

44. Yang, A. et al. MYC Inhibition Depletes Cancer Stem-like Cells in Triple-Negative Breast Cancer. Cancer Res. 77, 6641–6650 (2017).

45. Ramamoorthy, P., Dandawate, P., Jensen, R. A. & Anant, S. Celastrol and Triptolide Suppress Stemness in Triple Negative Breast Cancer: Notch as a Therapeutic Target for Stem Cells. Biomedicines 9, 482 (2021).

46. Li, J. et al. Triptolide-induced in vitro and in vivo cytotoxicity in human breast cancer stem cells and primary breast cancer cells. Oncol. Rep. 31, 2181–2186 (2014).

47. Das, B. & Kundu, C. N. Anti-Cancer Stem Cells Potentiality of an Anti-Malarial Agent Quinacrine: An Old Wine in a New Bottle. Anticancer Agents Med. Chem. 21, 416–427 (2021).

48. Nayak, D. et al. Quinacrine and curcumin synergistically increased the breast cancer stem cells death by inhibiting ABCG2 and modulating DNA damage repair pathway. Int. J. Biochem. Cell Biol. 119, 105682 (2020).

49. Das, B. et al. Quinacrine inhibits HIF-1α/VEGF-A mediated angiogenesis by disrupting the interaction between cMET and ABCG2 in patient-derived breast cancer stem cells. Phytomedicine Int. J. Phytother. Phytopharm. 117, 154914 (2023).

50. Cho, Y.-S., Kang, Y., Kim, K., Cha, Y.-J. & Cho, H.-S. The crystal structure of MPK38 in complex with OTSSP167, an orally administrative MELK selective inhibitor. Biochem. Biophys. Res. Commun. 447, 7–11 (2014).

51. Ganguly, R., Hong, C. S., Smith, L. G. F., Kornblum, H. I. & Nakano, I. Maternal Embryonic Leucine Zipper Kinase: Key Kinase for Stem Cell Phenotype in Glioma and Other Cancers. Mol. Cancer Ther. 13, 1393–1398 (2014).

52. Zhang, X. et al. MELK Inhibition Effectively Suppresses Growth of Glioblastoma and Cancer Stem-Like Cells by Blocking AKT and FOXM1 Pathways. Front. Oncol. 10, 608082 (2020).

53. Chung, S. et al. Development of an orally-administrative MELK-targeting inhibitor that suppresses the growth of various types of human cancer. Oncotarget 3, 1629–1640 (2012).

54. Spartinou, A., Nyktari, V. & Papaioannou, A. Granisetron: a review of pharmacokinetics and clinical experience in chemotherapy induced -nausea and vomiting. Expert Opin. Drug Metab. Toxicol. 13, 1289–1297 (2017).

55. Amini-Khoei, H. et al. Tropisetron suppresses colitis-associated cancer in a mouse model in the remission stage. Int. Immunopharmacol. 36, 9–16 (2016).

56. Pan, J. et al. Extracts of Zuo Jin Wan, a traditional Chinese medicine, phenocopies 5-HTR1D antagonist in attenuating Wnt/β-catenin signaling in colorectal cancer cells. BMC Complement. Altern. Med. 17, 506 (2017).

57. Brown, J. S. & Banerji, U. Maximising the potential of AKT inhibitors as anti-cancer treatments. Pharmacol. Ther. 172, 101–115 (2017).

58. Gallia, G. L. et al. Inhibition of Akt inhibits growth of glioblastoma and glioblastoma stem-like cells. Mol. Cancer Ther. 8, 386–393 (2009).

59. Eraslan, G., Simon, L. M., Mircea, M., Mueller, N. S. & Theis, F. J. Single-cell RNA-seq denoising using a deep count autoencoder. Nat. Commun. 10, 390 (2019).

60. Borgwardt, K. M. et al. Integrating structured biological data by Kernel Maximum Mean Discrepancy. Bioinforma. Oxf. Engl. 22, e49–57 (2006).

61. Paszke, A. et al. PyTorch: An Imperative Style, High-Performance Deep Learning Library. Preprint at https://doi.org/10.48550/arXiv.1912.01703 (2019).

62. Subramanian, A. et al. Gene set enrichment analysis: A knowledge-based approach for interpreting genome-wide expression profiles. Proc. Natl. Acad. Sci. U. S. A. 102, 15545–15550 (2005).

63. Napolitano, F. et al. gene2drug: a computational tool for pathway-based rational drug repositioning. Bioinformatics doi:10.1093/bioinformatics/btx800.

64. Liberzon, A. et al. The Molecular Signatures Database (MSigDB) hallmark gene set collection. Cell Syst. 1, 417–425 (2015).

65. Napolitano, F., Carrella, D., Gao, X. & di Bernardo, D. gep2pep: a bioconductor package for the creation and analysis of pathway-based expression profiles. Bioinformatics 36, 1944–1945 (2020).

