## supplementary text and figures for "AI identifies potent inducers of breast cancer stem cell differentiation based on adversarial learning from gene expression data"

Supplementary materials

#### Supplementary methods

##### The LINCS database

The Library of Integrated Network-based Cellular Signature (LINCS)[1] is a large collection of differential transcriptional profiles across multiple treated cell lines. The last release includes 118,050 GEPs across 41 cell lines of 2,169 perturbagens, 1,796 of which are small-molecule compounds. In this study, the level-5 data of the LINCS database were downloaded from the GEO website (GSE70138). CMAPPy[2] (version 4.0.1) was used to access the GCTX data format. The GEP signatures related to the 1,796 small-molecule treatments were used as the target domain dataset for the DREDDA model.

##### Data preparation and preprocessing

The gene expression counts from the hiPSC scRNA-seq dataset were distributed both sparsely and skewly, which may interfere with artificial neural network models convergence. Therefore, a zero-inflated negative binomial (ZINB) autoencoder model[3] was used for normalization and denoising. Since it is an unsupervised method, the ZINB-based model was trained using both the source and the target datasets and the estimated mean parameter ($\underline{M}$) of the model was used as the denoised version of the expression count matrix. The denoised count matrix was then transformed with the mapping $x \rightarrow log\,(x + \epsilon)$, where $\epsilon$ was set to $1.0 \times 10^{-5}$ to avoid undefined output values. Feature selection was performed on the source dataset by calculating the mutual information (MI) between each feature (gene) and the cluster labels. The top 1000 genes with the highest MI values were selected for subsequent analyses. LINCS level-5 data are already normalized. Only the genes covered by both of the datasets were selected as the final features and fed to the DREDDA model.

#### Description of the DREDDA architecture

On the high level, the DREDDA architecture is a three-module composite deep neural network (DNN) model consisting of (1) A domain-specific autoencoder; (2) A task classifier; (3) An adversarial domain classifier (se Figure 2 in the main text).

The domain-specific autoencoder is an autoencoder with two independent encoders and a shared decoder. Each of the two encoders processes profiles either from the source domain or the target domain. Here we denote them as $F_{enc\_s}(\,\cdot\,;\theta_{enc\_s})$ and $F_{enc\_t}(\,\cdot\,;\theta_{enc\_t})$, with $\theta_{enc\_s}$ and $\theta_{enc\_t}$ being their parameters, respectively. The use of two independent encoders yielded better performance as compared to a shared encoder (data not shown), likely due to the significant difference between the two input domains. With $X_s$ denoting a source domain input and $X_t$ denoting a target domain input, the two encoders then send the corresponding latent vectors $Z_s = F_{enc\_s}(X_s;\theta_{enc\_s})$ and $Z_t = F_{enc\_t}(X_t;\theta_{enc\_t})$ to the shared decoder, $F_{dec}(\,\cdot\,;\theta_{dec})$, for reconstruction. The reconstructed vectors $X_s'$ and $X_t'$ are used as inputs for the main task classifier and the adversarial task classifier.

The main task classifier is a multi-layer perceptron, here denoted as $F_{cls}(\,\cdot\,;\theta_{cls})$, estimating a classification probability for each of the four differentiation classes defined in the source domain, i.e. core, proliferative, early-primed and late-primed. For convenience, here we use $F_{cls,L}(X';\theta_{cls})$ to represent the intermediate network representation after the $L$-th layer of the task classifier when $X'$ is used as the input of the network.

The adversarial domain classifier, here denoted as $F_{adv}(\,\cdot\,;\theta_{adv})$ computes a binary decision on whether the reconstructed input comes from the source domain or the target domain.

Putting the three modules together, the DREDDA architecture aims to solve the so-called *un-supervised domain adaptation task*[4]. Indeed, while ground truth labels (i.e., the cell cluster labels) are only available for the source domain to train the model, the actual aim is to apply the trained model to the target domain (i.e., the LINCS dataset). Compared with Ganin et al.[4], the DREDDA model includes several modifications to make it suitable for gene expression data.

In further details, to train DREDDA on the unsupervised domain adaptation task, three objective (loss) functions were used. The first objective is the *task classification loss* for the classification task, which is a cross-entropy between prediction ($\hat{Y}$) ground truth label ($Y$, one-hot encoded):

$$L_{cls}^{(i)} = -\sum_{c}^{\#classes} Y_c^{(i)} \quad log\, \hat{Y}_c^{(i)} \tag{1}$$

where $\hat{Y}^{(i)}$ is the output vector of the task classifier $F_{cls}\left(X^{(i)\prime};\theta_{cls}\right)$ and $\hat{Y}_c^{(i)}$ takes its $c$-th dimension. The second objective is the *adversarial domain loss* for the adversarial domain classifier,

$$L_{adv}^{(i)} = -H^{(i)} \quad log\, \hat{H}^{(i)} - \left(1 - H^{(i)}\right) log\, (1 - \hat{H}^{(i)}) \tag{2}$$

where $H^{(i)} = 0$ if the $i$-th example belong to the source domain and $H^{(i)} = 1$ if it belongs to the target domain. $\hat{H}^{(i)}$ is the output of the adversarial domain classifier $F_{adv}(X^{(i)\prime};\theta_{adv})$. Inspired by the Deep Domain Confusion (DDC) framework[5], we further introduced a third objective, the *domain confusion loss*, which explicitly enforces similarity of intermediate network values between source domain and target domain examples by minimizing a Maximum Mean Discrepancy (MMD)[6] term,

$$\begin{aligned} MMD(\{source\},\{target\}) \\ = \| \frac{1}{\#source} \sum_{i \in \{source\}} \phi(X_i) \\ - \frac{1}{\#target} \sum_{j \in \{target\}} \phi(X_j) \| \end{aligned} \tag{3}$$

where $\phi(X_i)$ is the intermediate value in the network after it takes as input $X_i$. We chose $\phi(X_i)$ to represent the $L$-th intermediate layer of the task classifier, i.e., $\phi(X_i) = F_{cls,L}(X_i{}';\theta_{cls})$. Only the examples in one mini-batch are used to estimate MMD. The square of this term is used in the final objective function,

$$\begin{aligned} L = L_{cls} - L_{adv} + \lambda L_{dc} \\ = \left(\sum_{i \in \{source\}} L_{cls}^{(i)}\right) - \left(\sum_{i \in \{source\} \cup \{target\}} L_{adv}^{(i)}\right) \\ + \lambda MMD(\{source\},\{target\})^2 \end{aligned} \tag{4}$$

The sign in front of $L_{adv}$ is negative so as to maximize the domain adversarial classifier loss and thus discourage the use of domain-specific information during the training phase. The dependency of each loss term w.r.t. the model parameters can be inferred from the DREDDA architecture. Specifically, they are $L_{cls} = L_{cls}(\theta_{enc\_s}, \theta_{dec}, \theta_{cls})$, $L_{adv} = L_{adv}(\theta_{enc\_s}, \theta_{enc\_t}, \theta_{dec}, \theta_{adv})$ and $L_{dc} = L_{dc}(\theta_{enc\_s}, \theta_{enc\_t}, \theta_{dec}, \theta_{cls})$. The parameters are optimized via the following minimax objective,

$$L_{cls}\left(\theta_{enc\_s}, \theta_{dec}, \theta_{cls}\right) - L_{adv}\left(\theta_{enc\_s}, \theta_{enc\_t}, \theta_{dec}, \theta_{adv}\right) + L_{dc}(\theta_{enc\_s}, \theta_{enc\_t}, \theta_{dec}, \theta_{cls}) \quad (5)$$

#### Model Training and Implementation

Training of the network using the objective function described in Equation (5) can be done with a two-part update per training step. Specifically, one step updates the parameters that minimize the objective ($\theta_{enc\_s}, \theta_{enc\_t}, \theta_{dec}, \theta_{cls}$), while the other one updates the parameters that maximize the objective ($\theta_{adv}$). For each training step, an equal number of source domain and target domain examples are sampled from the source domain and target domain datasets. Using the sampled examples, $L_{cls}$ is computed using the sampled source domain examples, while $L_{adv}$ and $L_{dc}$ are computed using the sampled source and target domain examples. If plain minibatch stochastic gradient optimization is used, the parameters are updated as follows,

$$\theta_{enc\_s} \leftarrow \theta_{enc\_s} - \nabla_{\theta_{enc\_s}} L \theta_{enc\_t} \leftarrow \theta_{enc\_t} - \nabla_{\theta_{enc\_t}} L \quad (6)$$
$$\theta_{dec} \leftarrow \theta_{dec} - \nabla_{\theta_{dec}} L$$
$$\theta_{cls} \leftarrow \theta_{cls} - \nabla_{\theta_{cls}} L$$
$$\theta_{adv} \leftarrow \theta_{adv} + \nabla_{\theta_{adv}} L$$

Note that gradient *ascend* is performed on $\theta_{adv}$ while gradient *descent* is performed on all other parameters. The model is implemented using PyTorch 1.8[7] deep learning framework and can be run on any NVIDIA CUDA-capable GPUs with $\geq 10GB$ memory.

#### Model Testing and Predictions

After each training epoch, the model's performance was evaluated in the source domain based on multi-class classification loss and accuracy. At the same time, the adversarial classifier was evaluated on the full dataset based on the ability to discriminate between data points coming from the source domain and the target domain. The model was optimized towards highest source domain accuracy, with target domain accuracy within the 40%-60% range. The trained model was then applied to score each profile in the LINCS database and a final unique drug-related score was obtained by averaging over all the scores obtained for the same drug across different cell lines.

### Supplementary figures

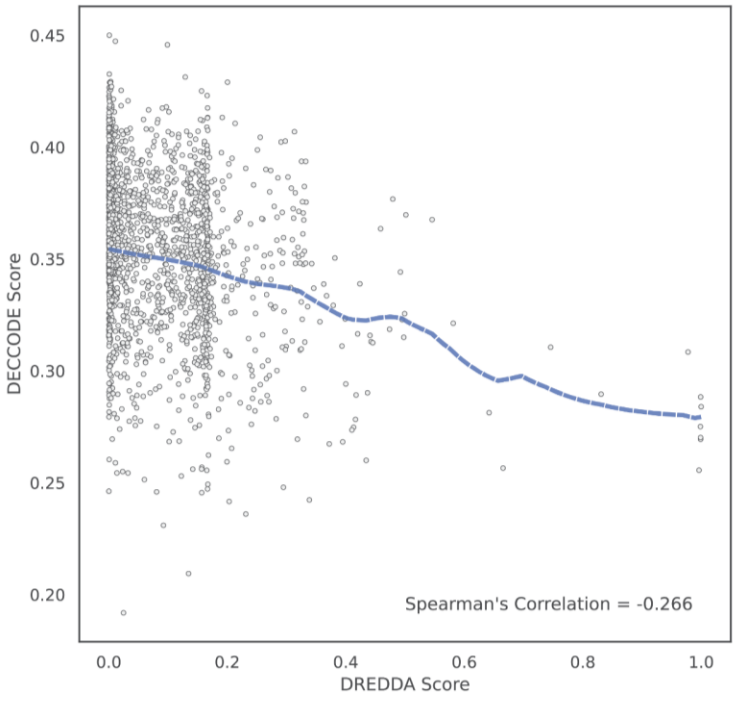


**Figure S1** Comparison between DREDDA scores (predicting differentiation) and DECCODE scores (predicting stemness). Although at low DREDDA scores there is no clear correlation between the two measures, high DREDDA scores tend to correspond to low DECCODE scores and high DECCODE scores tend to correspond to low DREDDA scores.

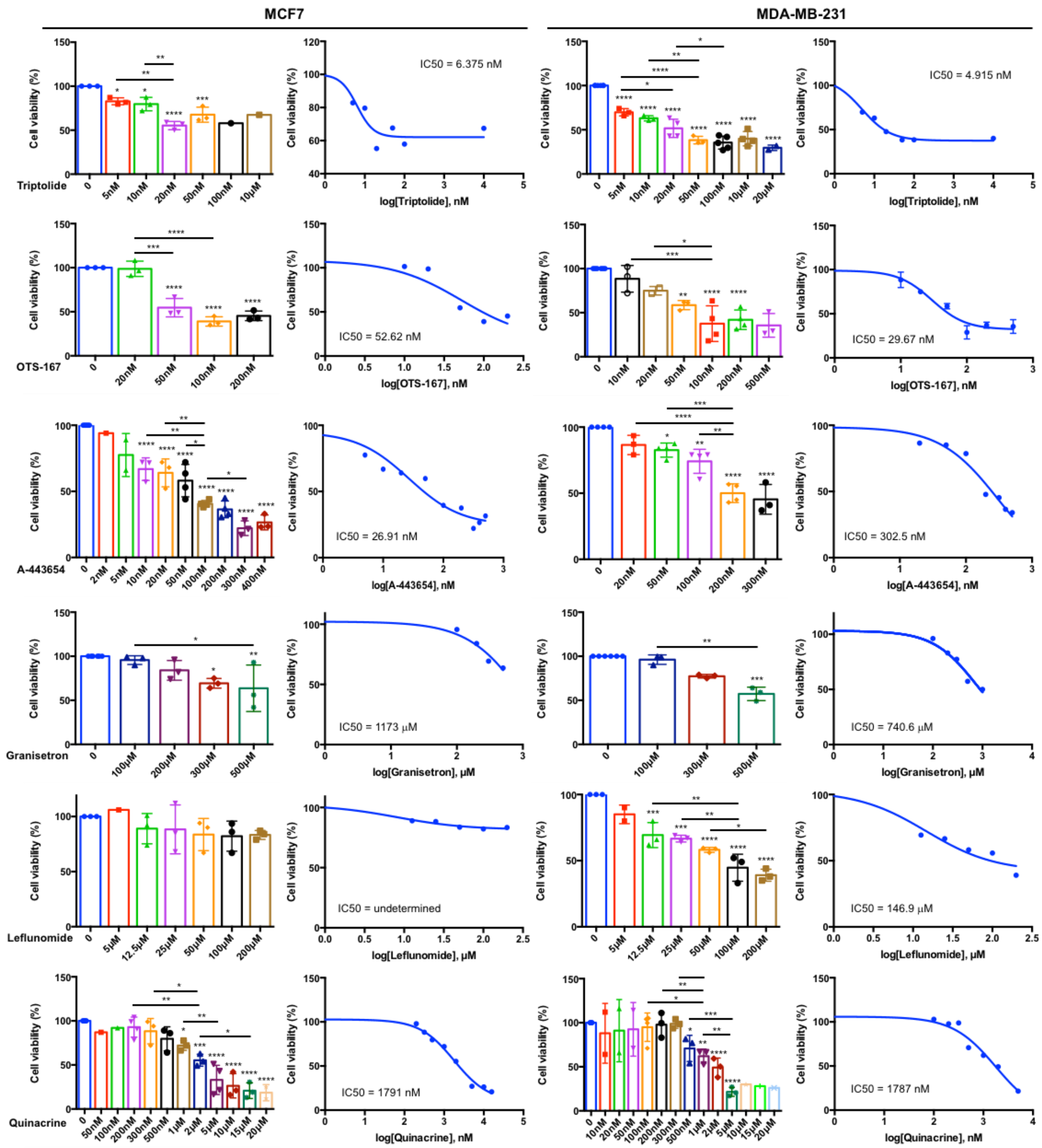


**Figure S2**. Percentage of cell viability and IC50 of MCF7 and MDA-MB-231 cells after exposure to increasing drug concentrations. MTT test results are reported as a percentage of cell viability. Each value was normalized with respect to its control. Dose-response curves were used to generate IC50. Statistics and IC50 curves were performed using Prism GraphPad 6.0. *, $P < 0.05$; **, $P < 0.01$; ***, $P < 0.001$; ****, $P < 0.0001$

### Supplementary tables

**Table S1** Top 30 molecules as prioritized by DREDDA.

| # | Drug | 2D structure | Functional Category | Notes |
|---|---|---|---|---|
| 1 | triptolide | 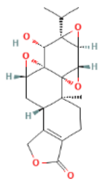 | Anti-proliferative agent | Has a role as an antispermatogenic agent and a plant metabolite. - ChEBI - http://www.ebi.ac.uk/chebi/searchId.do?chebiId=CHEBI:9747 |
| 2 | OTS-167 | 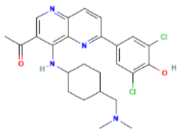 | MELK inhibitor | A naphthyridine derivative. - ChEBI - http://www.ebi.ac.uk/chebi/searchId.do?chebiId=CHEBI:95088 |
| 3 | CGP-60474 | 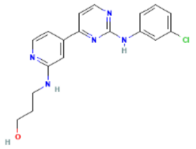 | CDK inhibitor | A substituted aniline. - ChEBI - http://www.ebi.ac.uk/chebi/searchId.do?chebiId=CHEBI:91339 |
| 4 | dinaciclib | 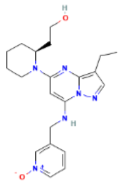 | CDK inhibitor | A pyrazolopyrimidine - ChEBI - http://www.ebi.ac.uk/chebi/searchId.do?chebiId=CHEBI:95060 |
| 5 | WZ-3105 | - | Small molecule kinase inhibitors | - |
| 6 | alvocidib | 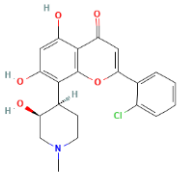 | CDK9 kinase inhibitor | A cyclin-dependent kinase 9 (CDK9) inhibitor, it has been studied for the treatment of acute myeloid leukaemia, arthritis and atherosclerotic plaque formation. It has a role as an antineoplastic agent, an EC 2.7.11.22 (cyclin-dependent kinase) inhibitor, an antirheumatic drug and an apoptosis inducer. - ChEBI - http://www.ebi.ac.uk/chebi/searchId.do?chebiId=CHEBI:47344 |

| # | Drug | 2D structure | Functional Category | Notes |
|---|---|---|---|---|
| 7 | BMS-387032 | 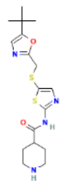 | CDK2 inhibitor | An ATP-competitive inhibitor of CDK2, CDK7 and CDK9 kinases and exhibits anti-cancer properties. It has a role as an apoptosis inducer, an antineoplastic agent, an EC 2.7.11.22 (cyclin-dependent kinase) inhibitor and an angiogenesis inhibitor.<br>- ChEBI -<br>http://www.ebi.ac.uk/chebi/searchId.do?chebiId=CHEBI:91399 |
| 8 | staurosporine | 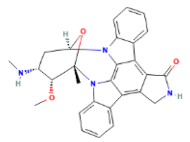 | General protein kinase inhibitor | Has a role as an EC 2.7.11.13 (protein kinase C) inhibitor, a geroprotector, a bacterial metabolite and an apoptosis inducer.<br>- ChEBI -<br>http://www.ebi.ac.uk/chebi/searchId.do?chebiId=CHEBI:15738 |
| 9 | AT-7519 | 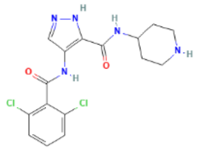 | CDK inhibitor | Has a role as an EC 2.7.11.22 (cyclin-dependent kinase) inhibitor and an antineoplastic agent.<br>- ChEBI -<br>http://www.ebi.ac.uk/chebi/searchId.do?chebiId=CHEBI:91326 |
| 10 | A-443654 | 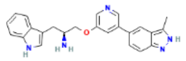 | Pan-Akt inhibitor | A member of indoles<br>- ChEBI -<br>http://www.ebi.ac.uk/chebi/searchId.do?chebiId=CHEBI:91351 |
| 11 | JNK-9L | 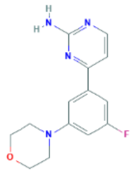 | JNK inhibitor | |
| 12 | mitoxantrone | 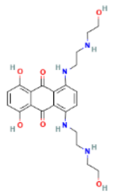 | TOPO II inhibitor | Has a role as an antineoplastic agent and an analgesic.<br>- ChEBI -<br>http://www.ebi.ac.uk/chebi/searchId.do?chebiId=CHEBI:50729 |
| 13 | Ro-4987655 | 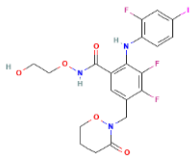 | MEK inhibitor | |

| # | Drug | 2D structure | Functional Category | Notes |
|---|---|---|---|---|
| 14 | binimetinib | 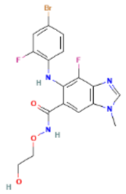 | MEK inhibitor | A MEK1 and MEK2 inhibitor (IC50= 12 nM). Approved by the FDA for the treatment of patients with unresectable or metastatic melanoma with a BRAF V600E or V600K mutation in combination with encorafenib. It has a role as an EC 2.7.11.24 (mitogen-activated protein kinase) inhibitor, an antineoplastic agent and an apoptosis inducer. - ChEBI - http://www.ebi.ac.uk/chebi/searchId.do?chebiId=CHEBI:145371 |
| 15 | R-547 | 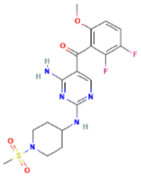 | CDK inhibitor | |
| 16 | bardoxolone-methyl | 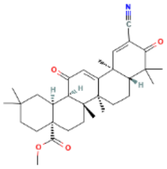 | NF-κB pathway inhibitor | Bardoxolone methyl is a member of cyclohexenones. - ChEBI - http://www.ebi.ac.uk/chebi/searchId.do?chebiId=CHEBI:177406 |
| 17 | PF-431396 | 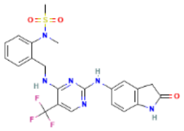 | FAK and PYK2 inhibitor | A sulfonamide. - ChEBI http://www.ebi.ac.uk/chebi/searchId.do?chebiId=CHEBI:91388 |
| 18 | camicinal | 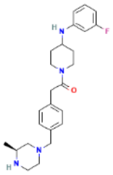 | Motilin agonist | Camicinal is a member of acetamides. - ChEBI - http://www.ebi.ac.uk/chebi/searchId.do?chebiId=CHEBI:177624 |
| 19 | AZD-5438 | 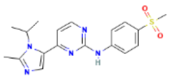 | CDK inhibitor | A sulfonamide. - ChEBI - http://www.ebi.ac.uk/chebi/searchId.do?chebiId=CHEBI:91419 |
| 20 | epirubicin | 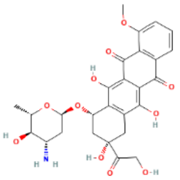 | DNA intercalating agents | Has a role as an EC 5.99.1.3 [DNA topoisomerase (ATP-hydrolysing)] inhibitor, an antineoplastic agent and an antimicrobial agent. - ChEBI - http://www.ebi.ac.uk/chebi/searchId.do?chebiId=CHEBI:47898 |

| # | Drug | 2D structure | Functional Category | Notes |
|---|---|---|---|---|
| 21 | AS-601245 | 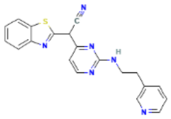 | JNK inhibitor | A member of benzothiazoles<br>- ChEBI -<br>http://www.ebi.ac.uk/chebi/searchId.do?chebiId=CHEBI:91345 |
| 22 | AZD-8330 | 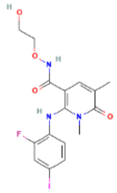 | MEK inhibitor | A pyridinecarboxamide. It is functionally related to a nicotinamide.<br>- ChEBI -<br>http://www.ebi.ac.uk/chebi/searchId.do?chebiId=CHEBI:91424 |
| 23 | leflunomide | 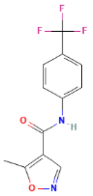 | Pyrimidine synthesis inhibitor | Has a role as a non-steroidal anti-inflammatory drug, an antineoplastic agent, an antiparasitic agent, an EC 1.3.98.1 [dihydroorotate oxidase (fumarate)] inhibitor, a hepatotoxic agent, a prodrug, a pyrimidine synthesis inhibitor, an immunosuppressive agent, an EC 3.1.3.16 (phosphoprotein phosphatase) inhibitor and a tyrosine kinase inhibitor.<br>- ChEBI -<br>http://www.ebi.ac.uk/chebi/searchId.do?chebiId=CHEBI:6402 |
| 24 | bruceantin | 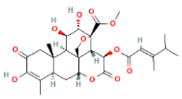 | Protein and DNA synthesis inhibitor | Bruceantin is a triterpenoid.<br>- ChEBI -<br>http://www.ebi.ac.uk/chebi/searchId.do?chebiId=CHEBI:3188 |
| 25 | SNX-2112 | 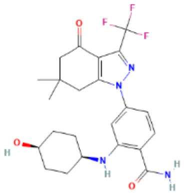 | Anti-proliferative agent | |
| 26 | PF-03758309 | 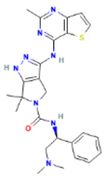 | PAK4 Inhibitor | An organic heterobicyclic compound, an organosulfur heterocyclic compound and an organonitrogen heterocyclic compound.<br>- ChEBI -<br>http://www.ebi.ac.uk/chebi/searchId.do?chebiId=CHEBI:93751 |
| 27 | rebastinib | 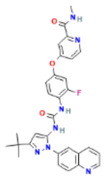 | TIE2 kinase inhibitor | Has a role as a tyrosine kinase inhibitor.<br>- ChEBI -<br>http://www.ebi.ac.uk/chebi/searchId.do?chebiId=CHEBI:62166 |

| # | Drug | 2D structure | Functional Category | Notes |
|---|---|---|---|---|
| 28 | granisetron | 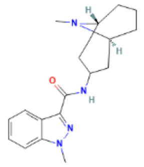 | Serotonin 5-HT3 receptor antagonist | A selective 5-HT3 receptor antagonist, it is used (generally as the monohydrochloride salt) to manage nausea and vomiting caused by cancer chemotherapy and radiotherapy, and to prevent and treat postoperative nausea and vomiting. It has a role as a serotonergic antagonist and an antiemetic. - ChEBI - http://www.ebi.ac.uk/chebi/searchId.do?chebiId=CHEBI:5537 |
| 29 | AEE-788 | 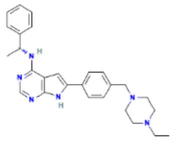 | RTK inhibitor | A potent inhibitor of human EGFR, VEGFR and HER2 receptor tyrosine kinases and exhibits anticancer and antiangiogenic activity. It has a role as an epidermal growth factor receptor antagonist, an EC 2.7.10.1 (receptor protein-tyrosine kinase) inhibitor, an antineoplastic agent, an angiogenesis inhibitor, a trypanocidal drug and an apoptosis inducer. - ChEBI - http://www.ebi.ac.uk/chebi/searchId.do?chebiId=CHEBI:40629 |
| 30 | deoxycholic-acid | 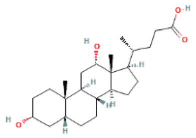 | Oxidative agent | A bile acid, a dihydroxy-5beta-cholanic acid and a C24-steroid. It is a conjugate acid of a deoxycholate. - ChEBI - http://www.ebi.ac.uk/chebi/searchId.do?chebiId=CHEBI:28834 |

**Table S2** Top-enriched GO terms under the Biological Process category resulting from drug set enrichment analysis (DSEA) of the top 30 drugs vs bottom 30 drugs

| Pathway | ES | PVadj |
|---|---|---|
| Morphogenesis of a polarized epithelium | -0.97 | 1.11E-10 |
| Nuclear envelope organization | -0.97 | 1.11E-10 |
| Female meiosis I | -0.97 | 1.11E-10 |
| Dosage compensation | -0.97 | 1.11E-10 |
| Pyrimidine containing compound salvage | -0.97 | 1.11E-10 |
| Protein import | -0.97 | 1.11E-10 |
| RNA 3’ end processing | -0.97 | 1.11E-10 |
| MRNA 3’ end processing | -0.97 | 1.11E-10 |
| Positive regulation of type I interferon production | -0.97 | 1.11E-10 |
| RNA stabilization | -0.97 | 1.11E-10 |
| Cell cycle G2-M phase transition | -0.97 | 1.11E-10 |

| | | |
|---|---|---|
| tRNA transport | -0.97 | 1.11E-10 |
| Nuclear transport | -0.97 | 1.11E-10 |
| Telomerase RNA localization | -0.97 | 1.11E-10 |
| ncRNA export from nucleus | -0.97 | 1.11E-10 |
| Regulation of protein localization to nucleus | -0.97 | 1.11E-10 |
| Regulation of signal transduction by p53 class mediator | -0.97 | 1.11E-10 |
| Positive regulation of cyclin dependent protein kinase activity | -0.97 | 1.11E-10 |
| Regulation of protein localization to chromosome telomeric region | -0.97 | 1.11E-10 |
| Positive regulation of telomerase rna localization to cajal body | -0.97 | 1.11E-10 |
| Photoreceptor cell maintenance | 0.97 | 1.11E-10 |
| Pattern specification process | 0.93 | 3.00E-09 |
| Hormone transport | 0.93 | 3.00E-09 |
| Monovalent inorganic cation transport | 0.93 | 3.00E-09 |
| Hormone metabolic process | 0.93 | 3.00E-09 |
| Activation of janus kinase activity | 0.93 | 3.00E-09 |
| Photoreceptor cell differentiation | 0.93 | 3.00E-09 |
| Regulation of calcium ion transmembrane transport | 0.93 | 3.00E-09 |
| Regulation of neurotransmitter levels | 0.9 | 5.79E-08 |
| Eye photoreceptor cell differentiation | 0.9 | 5.79E-08 |
| Regionalization | 0.9 | 5.79E-08 |
| Regulation of systemic arterial blood pressure | 0.9 | 5.79E-08 |
| Potassium ion transport | 0.9 | 5.79E-08 |
| Excretion | 0.9 | 5.79E-08 |
| Regulation of blood pressure | 0.9 | 5.79E-08 |
| Specification of symmetry | 0.9 | 5.79E-08 |
| Regulation of hormone levels | 0.9 | 5.79E-08 |
| Response to auditory stimulus | 0.9 | 5.79E-08 |
| Organic anion transport | 0.9 | 5.79E-08 |
| Organic hydroxy compound transport | 0.9 | 5.79E-08 |

**Table S3** Top-enriched GO terms under the Cellular Component category resulting from drug set enrichment analysis (DSEA) of the top 30 drugs vs bottom 30 drugs

| **Pathway** | **ES** | **PVadj** |
|---|---|---|
| Nuclear periphery | -1.00 | 0.00E+00 |

| | | |
|---|---|---|
| Nuclear ubiquitin ligase complex | -0.97 | 1.11E-12 |
| Fibrillar center | -0.97 | 1.11E-12 |
| Nuclear matrix | -0.97 | 1.11E-12 |
| Aminoacyl trna synthetase multienzyme complex | -0.97 | 1.11E-12 |
| Protein acetyltransferase complex | -0.97 | 1.11E-12 |
| Nuclear membrane | -0.97 | 1.11E-12 |
| Histone deacetylase complex | -0.93 | 3.00E-11 |
| Commitment complex | -0.93 | 3.00E-11 |
| Transcription export complex | -0.93 | 3.00E-11 |
| Chromosome centromeric region | -0.93 | 3.00E-11 |
| Kinetochore | -0.93 | 3.00E-11 |
| Condensed chromosome centromeric region | -0.93 | 3.00E-11 |
| Spindle pole | -0.93 | 3.00E-11 |
| Nuclear envelope | -0.93 | 3.00E-11 |
| Rna polymerase III complex | -0.93 | 3.00E-11 |
| Spliceosomal complex | -0.93 | 3.00E-11 |
| U1 snRNP | -0.93 | 3.00E-11 |
| Centrosome | -0.93 | 3.00E-11 |
| Spindle | -0.93 | 3.00E-11 |
| Cilium | 0.93 | 3.00E-11 |
| Photoreceptor outer segment | 0.90 | 5.79E-10 |
| Intrinsic component of the cytoplasmic side of the plasma membrane | 0.90 | 5.79E-10 |
| Photoreceptor connecting cilium | 0.90 | 5.79E-10 |
| Interphotoreceptor matrix | 0.90 | 5.79E-10 |
| Ciliary transition zone | 0.90 | 5.79E-10 |
| Photoreceptor outer segment membrane | 0.90 | 5.79E-10 |
| Inhibitory synapse | 0.90 | 5.79E-10 |
| Ciliary membrane | 0.90 | 5.79E-10 |
| Ciliary plasm | 0.90 | 5.79E-10 |
| 9plus0 non motile cilium | 0.90 | 5.79E-10 |
| Voltage gated sodium channel complex | 0.87 | 8.25E-09 |
| Photoreceptor inner segment | 0.87 | 8.25E-09 |
| Axonemal dynein complex | 0.87 | 8.25E-09 |
| Cation channel complex | 0.87 | 8.25E-09 |
| Potassium channel complex | 0.87 | 8.25E-09 |
| Sodium channel complex | 0.87 | 8.25E-09 |
| Sperm midpiece | 0.87 | 8.25E-09 |
| Non motile cilium | 0.87 | 8.25E-09 |
| Photoreceptor cell cilium | 0.87 | 8.25E-09 |

**Table S4**. Molecules, IC50, and working concentrations for the CSC targeting experiments.

| Molecule Name | IDs | MCF7 concentrations | MDA-MB-231 concentrations |
|---|---|---|---|
| triptolide | BRD-K81258678<br>Sigma (cat. # S-645900) | IC50 = 6.375 nM<br>10nM; 20nM | IC50 = 4.915 nM<br>10nM; 50nM |
| OTS-167 | BRD-K53417444<br>Vinci-Biochem (cat. # CAY-16873-5) | IC50 = 52.62 nM<br>50nM; 100nM | IC50 = 29.67 nM<br>50nM; 100nM |
| granisetron | BRD-A10967948<br>DBA (cat. # SC-203983) | IC50 = 1173 µM<br>300µM; 500µM | IC50 = 740.6 µM<br>300µM; 500µM |
| A-443654 | BRD-K88573743<br>DBA (cat. # HY-10425) | IC50 = 26,91 nM<br>50nM; 100nM | IC50 = 302.5 nM<br>50nM; 100nM |
| leflunomide | BRD-K78692225<br>DBA (cat. # HY-B0083) | IC50 = Undetermined<br>100µM; 200µM | IC50 = 146.9 µM<br>25µM; 50µM |
| quinacrine | BRD-A45889380<br>DBA (cat. # HY-13735A) | IC50 = 1.791 µM<br>500nM; 5µM | IC50 = 1.787 µM<br>500nM; 5µM |

**Table S5.** Detailed Network Architecture of DREDDA

| Network Module | Architecture |
|---|---|
| Source domain encoder | Input dim: (760,) |
| | Fully Connected Layer: (760,) ➡(100,) |
| | Fully Connected Layer: (100,) ➡(50,) |
| Target domain encoder | Input dim: (760,) |
| | Fully Connected Layer: (760,) ➡(100,) |
| | Fully Connected Layer: (100,) ➡(50,) |
| Shared decoder | Fully Connected Layer: (50,) ➡(100,) |
| | Fully Connected Layer: (100,) ➡(760,) |
| Task classifier | Fully Connected Layer: (760,) ➡(100,) |
| | Fully Connected Layer: (100,) ➡(80,) |
| | Fully Connected Layer: (80,) ➡(60,) |
| | Fully Connected Layer: (60,) ➡(40,) |
| | Fully Connected Layer: (40,) ➡(20,) |
| | Task Output Layer: (20,) ➡(4,) |

| | |
|---|---|
| Adversarial domain classifier | Fully Connected Layer: (60,) ➡(40,) |
| | Adversarial Output Layer: (40,) ➡(2,) |
